# A widely distributed gene cluster compensates for uricase loss in hominids

**DOI:** 10.1101/2022.07.24.501321

**Authors:** Yuanyuan Liu, J. Bryce Jarman, Yen S. Low, Steven Huang, Haoqing Chen, Mary E. DeFeo, Kazuma Sekiba, Bi-Huei Hou, Calyani Ganesan, Alan C. Pao, Saurabh Gombar, Dylan Dodd

## Abstract

Approximately 15% of US adults have circulating levels of uric acid above its solubility limit, which is causally linked to the disease gout. In most mammals, uric acid elimination is facilitated by the enzyme uricase. However, human uricase is a pseudogene, having been inactivated early in hominid evolution. Though it has long been known that uric acid is eliminated in the gut, the role of the gut microbiota in hyperuricemia has not been studied. Here we identify a widely distributed bacterial gene cluster that encodes a pathway for uric acid degradation. Stable isotope tracing demonstrates that gut bacteria metabolize uric acid to xanthine or short chain fatty acids. Ablation of the microbiota in uricase-deficient mice causes severe hyperuricemia, and anaerobe-targeted antibiotics increase the risk of gout in humans. These data reveal a role for the gut microbiota in uric acid excretion and highlight the potential for microbiome-targeted therapeutics in hyperuricemia.

## Introduction

Uric acid is an intermediate in purine degradation in mammals. In most mammals, uric acid is converted to freely soluble allantoin via urate oxidase (uricase) which is then excreted via the kidney. However, early in hominid evolution progressive mutations occurred in the uricase gene decreasing its activity until uricase function was completely lost (Kratzer et al., 2014). Although uricase pseudogenization may have been beneficial for our ancestors (Chang, 2014; Johnson et al., 2005; Kratzer et al., 2014), in modern times it has become a liability. Approximately 14.6% of the US population has hyperuricemia (defined by plasma levels of uric acid > 6.8 mg/dL (> 0.4 mM)) and 3.9% have clinical features of gout, a painful inflammatory arthritis caused by precipitation of uric acid crystals (Singh et al., 2019). Therapies for gout include inhibitors of xanthine oxidase – upstream of uric acid in the purine metabolism pathway – or drugs that block reabsorption of uric acid in the proximal renal tubule. Most of these medications suffer either from poor efficacy, poor compliance, or intolerable side effects, thus new therapies for gout are needed.

Two independent lines of evidence suggest that the gut is an important site for uric acid elimination in humans: First, radioisotope studies in healthy individuals revealed that ∼1/3 of uric acid is disposed from the gut; in patients with kidney disease, this proportion rises to ∼2/3 (**Figure 1A**) (Sorensen, 1965). Second, variant alleles in the intestinal/renal transporter ABCG2 diminish intestinal uric acid elimination (Hoque et al., 2020) and are risk factors for hyperuricemia and gout (Kottgen et al., 2013; Tin et al., 2019). While it is presumed that bacteria in the gut break down uric acid to products that are absorbed and excreted by the host (Sorensen, 1965), uric acid metabolism by commensal gut bacteria has not been studied. Intriguingly, recent data in uricase-deficient mice (Pierzynowska et al., 2020) and in humans with hyperuricemia and normal to Stage 2 chronic kidney disease (estimated glomerular filtration > 60 mL/min) (Terkeltaub et al., 2022) suggest that enzymatic degradation of uric acid within the gut can reduce serum uric acid levels. Collectively, these observations indicate that uric acid metabolism by gut bacteria could be important for maintaining low serum uric acid levels.

**Figure 1.**
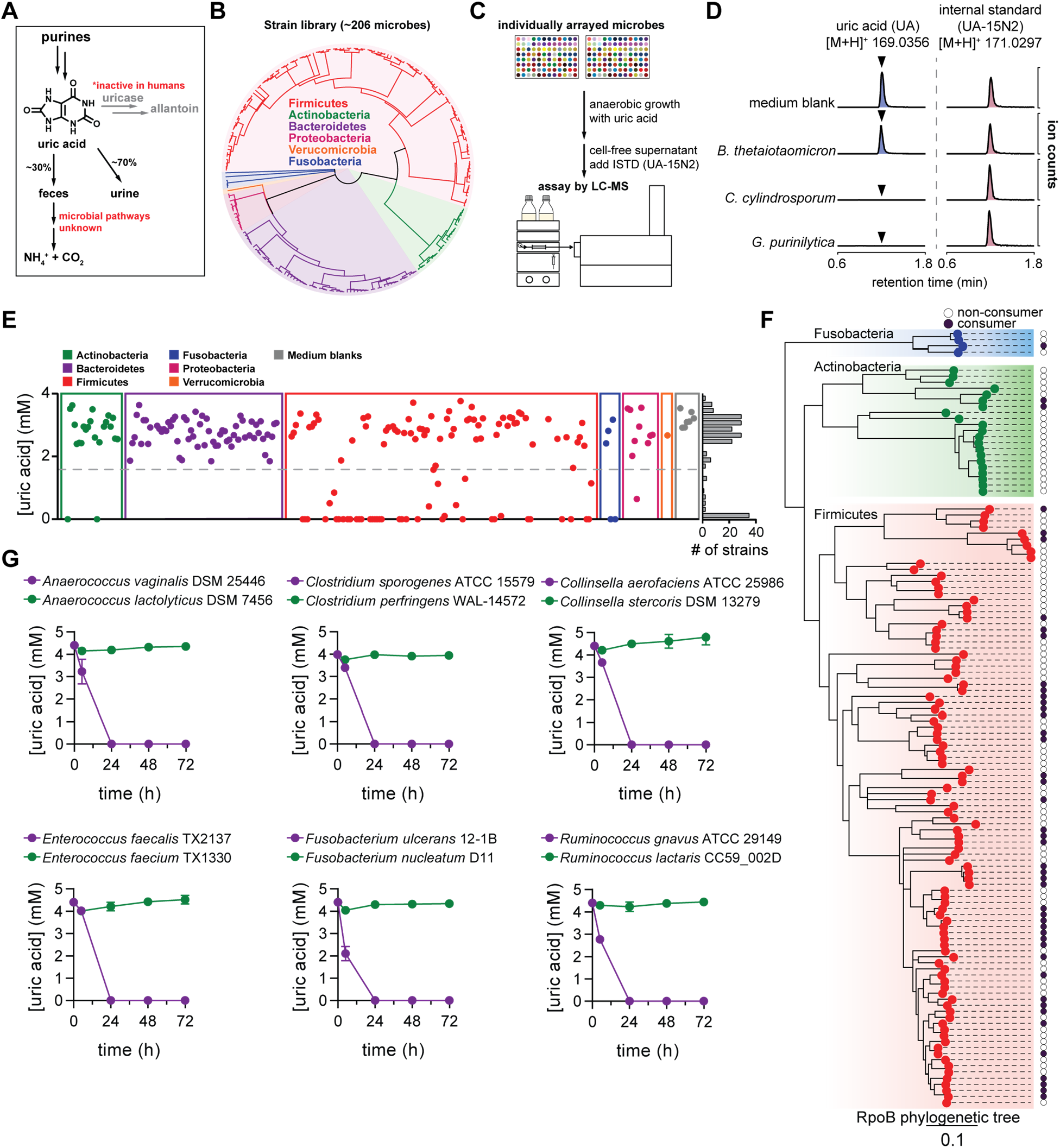
Anaerobic uric acid metabolism is widespread among human gut bacteria. A) Overview of purine metabolism in humans. Uricase is a pseudogene and ∼2/3 of uric acid is eliminated in urine, with an additional 1/3 being excreted in the intestine. B) Phylogenetic distribution of human gut bacteria in the strain library used for this study. C) Overview of experimental approach to screen for uric acid metabolism. Bacteria were cultured in rich medium with uric acid and uric acid levels in cell-free supernatants were quantified after 48 h by isotope-dilution LC-MS. D) Extracted ion chromatograms for uric acid and the uric acid internal standard (ISTD; [^15^N_2_]-uric acid) in medium blank and after incubation with a non-consumer (*B. thetaiotaomicron*) and two known obligate uric acid consuming bacteria (*C. cylindrosporum* and *G. purinilytica*). E) Results from uric acid screen in rich medium, grouped by phylum. Each dot represents a single bacterial strain. The frequency of strains is shown on the right of the plot. F) Phylogenetic distribution of uric acid consuming bacteria within the Actinobacteria, Fusobacteria, and Firmicutes phyla. Dark purple dots represent strains that consume >50% of the uric acid. Only those strains for which assembled genomes are available were included. G) Uric acid consumption in closely related bacteria during growth in rich media. For D and E, data represent the results from a single experiment. For G, data represent the means ± standard deviations of n = 3 biological replicates.

Here, we report that many gut bacteria consume uric acid anaerobically, converting it to either xanthine or lactate and the short chain fatty acids (SCFAs) acetate and butyrate. Transcriptional profiling and genetics reveal a gene cluster that is required for conversion of uric acid to lactate and SCFAs and is widely distributed across phylogenetically distant bacterial taxa. We find that human gut bacteria compensate for the loss of uricase in a mouse model, and that antibiotics targeting anaerobic bacteria, which would ablate gut bacteria, increase the risk for developing gout in humans. Together, our findings uncover a previously unknown mechanism by which gut bacteria contribute to uric acid homeostasis in the host.

## Results

### Anaerobic uric acid metabolism is widespread among gut bacteria

While uric acid metabolism is well known to occur among aerobic bacteria, anaerobic uric acid metabolism has been described in only a few obligate purine-degrading bacteria isolated from soil. Early biochemical studies with *Gottschalkia acidurici, G. purinilytica* and *Clostridium cylindrosporum*, all soil isolates, established the enzymatic activities involved in anaerobic uric acid metabolism; however, the identity of genes supporting this purinolytic pathway is not known (Hartwich et al., 2012; Poehlein et al., 2015a; Poehlein et al., 2015b). Thus, no marker genes are available to query gut bacterial genomes for uric acid metabolism.

To identify uric acid consuming gut bacteria, we assembled a phylogenetically diverse library consisting of 206 human gut isolates representing 6 phyla and 38 bacterial families (**Figure 1B**). We cultured these strains anaerobically for 48 h in rich media supplemented with uric acid then quantified remaining uric acid by isotope dilution LC-MS (**Figure 1C**). We validated the assay using two known uric acid degrading bacteria (*C. cylindrosporum* and *G. purinilytica*) (Durre and Andreesen, 1983) and a well-characterized commensal gut bacterium, *Bacteroides thetaiotaomicron*.

While chromatographic peaks for uric acid were comparable between the medium control and *B. thetaiotaomicron* supernatants, peaks were absent for *C. cylindrosporum* and *G. purinilytica*, demonstrating these two bacteria consume uric acid (**Figure 1D**). By screening the entire library, we found that uric acid consumption was remarkably widespread with over 1/5 of bacteria (47/206) consuming >50% of uric acid after 48 h of anaerobic growth (**Figure 1E** and **Table S1**). Uric acid consumption was distributed across 4 phyla (Actinobacteria, Firmicutes, Fusobacteria, and Proteobacteria), but notably absent in Bacteroidetes. We repeated this screen with an expanded library of strains under carbohydrate limiting conditions (**Figure S1A** and **Table S2**). Results from the second screen: i) confirmed findings for most of the organisms in the first screen, ii) identified additional uric acid consuming strains, bringing the total to 60/243 strains tested (not including the soil isolates *C. cylindrosporum* and *G. purinilytica*), and iii) suggested that some strains consume more uric acid in the absence of carbohydrates (**Figure S1B**). This latter observation implies that nutrient conditions influence the capacity of gut bacteria to consume uric acid and our screen may underestimate the number of uric acid consuming bacteria.

By combining results from the two screens, we found that the capacity for uric acid consumption varies widely, even among closely related bacteria (**Figure 1F**). Notable examples of this disconnect between phylogeny and physiology occur within the *Anaerococcus, Clostridium, Collinsella, Enterococcus, Fusobacterium, Streptococcus, Blautia, Dorea*, and *Ruminococcus* genera (**Figure 1F**). By culturing a subset of these related species with uric acid, we confirmed that uric acid metabolism is not strictly conserved across phylogenetic lineages (**Figure 1G** and **Table S3**). These findings suggest that the capacity to consume uric acid may have been gained or lost multiple times during bacterial evolution.

### Gut bacteria convert uric acid into xanthine and short chain fatty acids

Having identified a large number of uric acid consuming gut bacteria, we next asked what is the metabolic fate of uric acid in these bacteria. Given that xanthine is a common intermediate in purine metabolism, we reasoned that gut bacteria might convert uric acid to xanthine. To test this, we quantified xanthine in the supernatants of bacteria cultured with uric acid in our two screens. Among the bacteria that consumed more than 50% of uric acid in the medium, we found that some also accumulated xanthine in supernatants (**Figure S2, Table S1, Table S2**). However, these xanthine producing strains represented just a subset of uric acid consumers, indicating that many bacteria produce other yet unidentified metabolites.

We reasoned that bacteria might break uric acid down to metabolic end products such as short chain fatty acids (SCFAs). To address this, we synthesized uniformly labeled [^13^C_5_]-uric acid where each of the five carbons was labeled and performed stable isotope tracing in xanthine-producing and non-producing strains (**Figure 2A**). First, we compared the metabolic fate of [^13^C_5_]-uric acid in cultures of a xanthine producing strain (*Blautia sp*. KLE 1732) and a xanthine non-producing strain (*Clostridium sporogenes* ATCC 15579). Over time, *Blautia sp*. KLE 1732 consumed [^13^C_5_]-uric acid and the M+5 isotope of xanthine accumulated in culture supernatants (**Figure 2B**). In contrast, while [^13^C_5_]-uric acid was depleted in *C. sporogenes* ATCC 15579 cultures, M+5 xanthine did not accumulate (**Figure 2C**). Rather, we detected an increase in the M+2 isotopes of acetate and butyrate, suggesting that *C. sporogenes* converts uric acid to SCFAs (**Figure 2C**). M+2 acetate peaks also appeared in the *Blautia sp*. KLE 1732 cultures, but the levels were equivalent in both unlabeled and labeled uric acid supplemented cultures (**Figure 2B** and **Table S4**), indicating that these peaks represent natural M+2 isotope abundance arising from acetate produced during growth (**Figure S3** and **Table S4**). Therefore, for all strains tested, we subtracted isotopes during growth with unlabeled uric acid from isotopes during growth with labeled uric acid. These experiments revealed that certain strains of bacteria (e.g., *Blautia sp*. KLE 1732 and *Coprococcus comes* ATCC 27758) convert uric acid to xanthine without breaking xanthine down further (**Figure 2D-E** and **Table S4**). However, other bacteria break uric acid down completely, yielding lactate and the SCFAs, acetate and butyrate (**Figure 2D-E** and **Table S4**).

**Figure 2.**
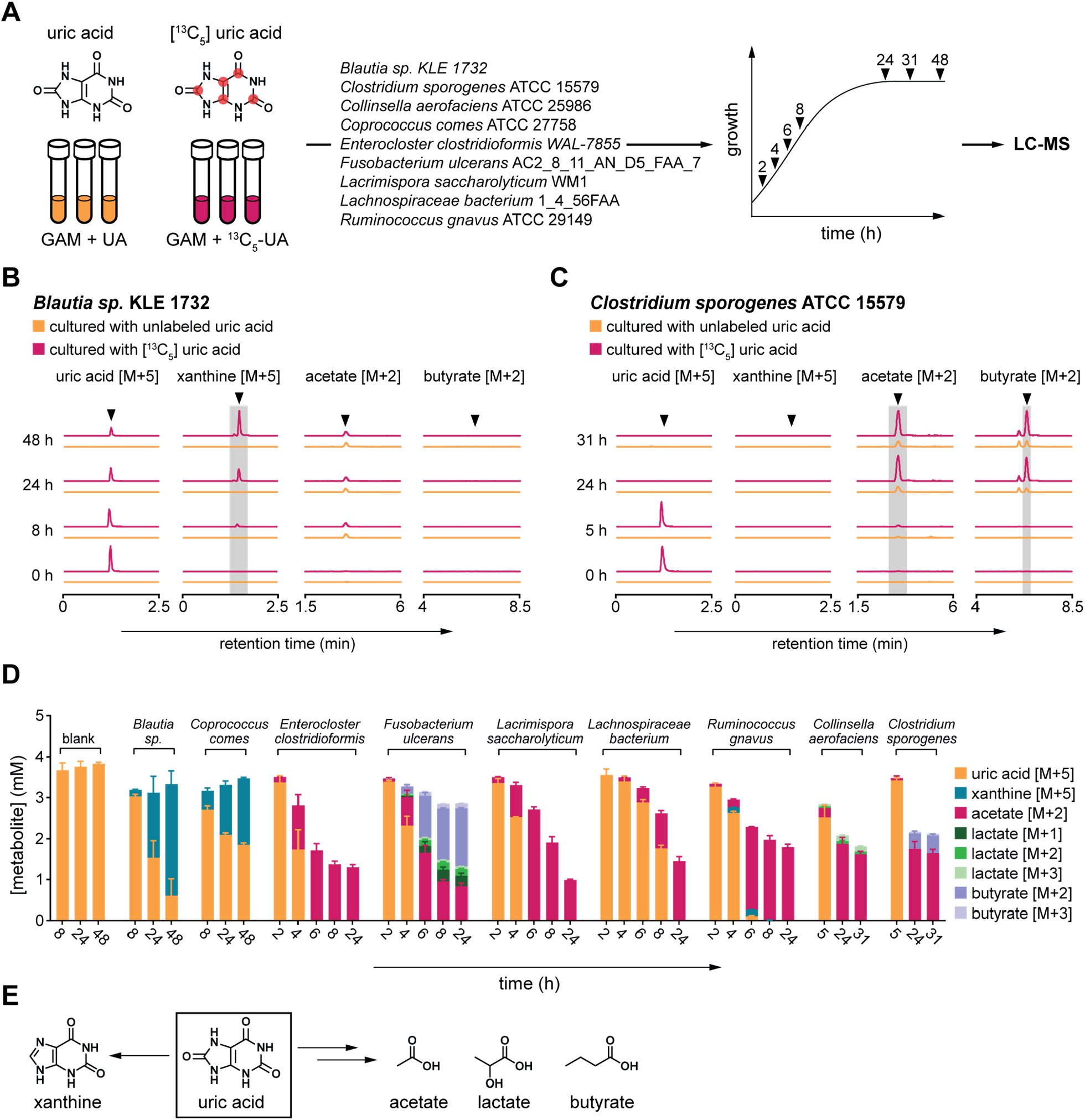
Gut bacteria convert uric acid into xanthine or lactate and short chain fatty acids. A) Overview of stable isotope tracing. Bacteria were cultured in rich media containing either unlabeled or uniformly labeled [^13^C_5_] uric acid and metabolites were quantified at indicated times by LC-MS. B) Extracted ion chromatograms for labeled substrates or products when *Blautia sp*. KLE 1732 was cultured with labeled or unlabeled uric acid. C) Extracted ion chromatograms for labeled substrates or products when *Clostridium sporogenes* ATCC 15579 was cultured with labeled or unlabeled uric acid. D) Labeled substrates and products detected in cell-free culture supernatants of all nine bacteria studied. To account for natural isotope abundances, labeled metabolites detected in unlabeled cultures were subtracted from labeled metabolites detected in labeled cultures. E) Uric acid is converted either to xanthine or lactate and the short chain fatty acids acetate and butyrate. For B and C, arrows indicate expected retention times for indicated compounds. For C, the peak eluting 0.3 min before butyrate [M+2] was identified as isobutyrate [M+2]. For B and C, experiments were performed in triplicate and representative data are shown. For D, data represent the means ± standard deviations of n = 3 biological replicates.

### Identification of a uric acid-inducible gene cluster required for anaerobic uric acid metabolism

To provide insights into which genes might be involved in conversion of uric acid to short chain fatty acids, we cultured three organisms (*C. sporogenes, L. saccharolyticum*, and *C. aerofaciens*) in rich medium with or without uric acid and performed RNAseq analysis (**Figure 3A**). These organisms were selected to represent phylogenetically distinct bacteria (two Firmicutes and one Actinobacteria) that convert uric acid to SCFAs (**Figure 2D**). Analysis of the RNA-seq data revealed that most of the uric acid inducible genes were not shared across these three bacteria, revealing an organism-specific response to uric acid availability (**Figure 3B**). However, we found 5 genes (*ygeX, ygeY, ygeW, ygfK*, and *ssnA*) that were shared across the three bacteria (**Figure 3B**) and were among the most highly induced genes in each of these organisms (**Figure 3C** and **Tables S5-7**). We then analyzed the genomic context for these genes and found that they map to a discrete gene cluster shared across the three bacteria (**Figure 3D**). Notably, annotations for these genes derive from *Escherichia coli* where they code for enzymes whose activities are predicted at the family level, but for which substrate specificities and cellular roles are unknown. Putative annotations for these gene products include ammonia lyase (YgeX), peptidase (YgeY), carbamoyl transferase (YgeW), oxidoreductase (YgfK), and aminohydrolase (SsnA) which are enzymes that may reduce and cleave bonds present in uric acid. These findings reveal a conserved set of uric acid-inducible genes that are shared across phylogenetically distinct gut bacteria.

**Figure 3.**
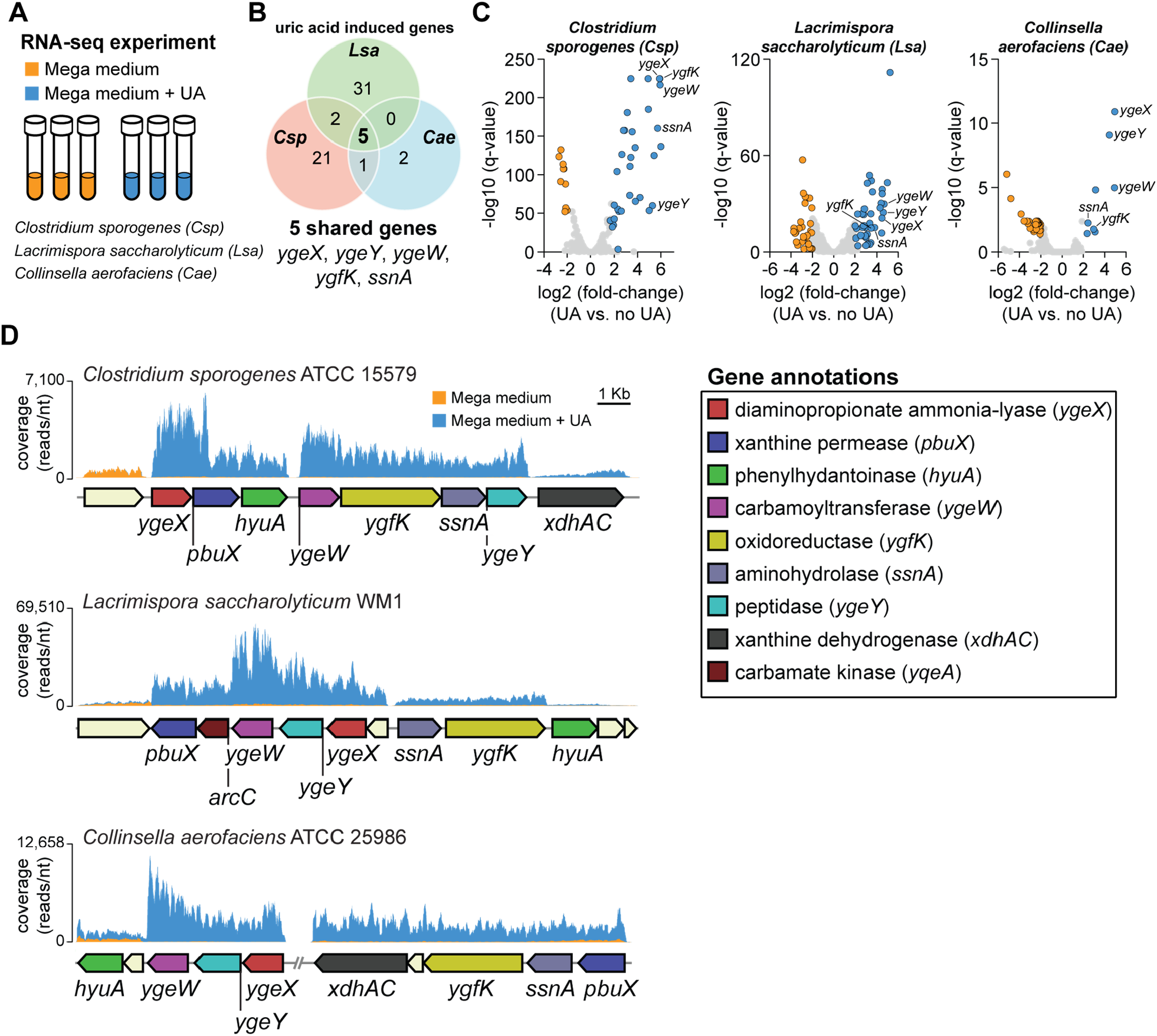
RNA-seq reveals a novel uric acid-inducible gene cluster in gut bacteria. A) Overview of experimental design. Three organisms (*C. sporogenes, L. saccharolyticum*, and *C. aerofaciens*) were cultured in rich medium with and without supplemental uric acid and transcriptomes were analyzed by RNA-seq. B) Venn diagram showing significantly induced genes for all three organisms (FDR corrected *P*-value (q-value) < 0.05, fold-change > 4). C) Volcano plots showing differentially regulated genes in the three organisms. Cut-offs include FDR corrected *P*-value (q-value) < 0.05 and |fold-change| > 4. Each dot represents a single gene. Blue dots represent genes that are induced when uric acid is present and orange dots represent genes that are repressed when uric acid is present. D) Genomic context and RNA-seq coverage for conserved uric acid inducible genes. Genes are color coded according to gene annotations. For RNA-seq experiments, three biological replicates were performed for each condition. For D, representative data are shown.

To test whether these genes are involved in uric acid metabolism, we individually disrupted each of the *C. sporogenes* genes using ClosTron (Heap et al., 2010) (**Figure 4A**). We then used stable isotope tracing to quantify uric acid metabolism and conversion to SCFAs by wild-type and mutant *C. sporogenes*. Consistent with our previous results, the wild-type *C. sporogenes* strain consumed all the uric acid, converting it to a mixture of M+2 acetate, M+2 butyrate, and trace amounts of M+3 butyrate (**Figure 4B** and **Table S8**). By comparison, the mutant strains were each either partially (*ygeX, pbuX, hyuA, ygeW, ygfK, ssnA, ygeY*) or completely (*xdhAC*) blocked in uric acid metabolism (**Figure 4B** and **Table S8**). Notably, neither labeled acetate nor labeled butyrate were detected in culture supernatants of any of the mutant strains (**Figure 4B** and **Table S8**). These findings provide evidence that the uric acid-inducible genes in *C. sporogenes* are required for conversion of uric acid to SCFAs including acetate and butyrate.

**Figure 4.**
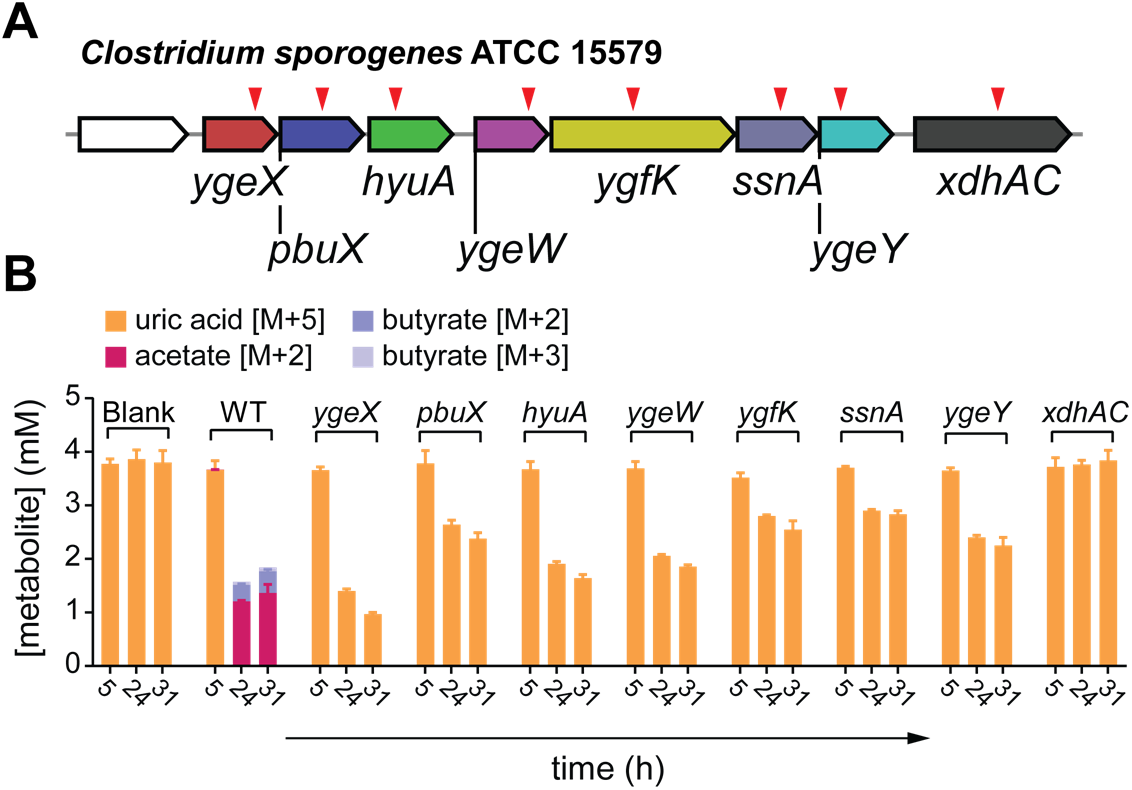
Uric acid inducible genes are required for conversion of uric acid to short chain fatty acids. A) Individual mutants (indicated by red triangles) were generated in the *C. sporogenes* uric acid inducible gene cluster using the ClosTron system. B) Stable isotope tracing in wild-type and mutant *C. sporogenes* strains. Strains were cultured in rich medium containing either labeled or unlabeled uric acid. Labeled substrates and products in cell-free supernatants were quantified by LC-MS. To account for natural isotope abundances, labeled metabolites detected in unlabeled cultures were subtracted from labeled metabolites detected in labeled cultures. For B, data represent the means ± standard deviations of n = 3 biological replicates.

### Uric acid-inducible genes are widely distributed across human gut bacteria

To evaluate whether other uric acid consuming strains in our library harbor this gene cluster, we performed BLASTp searches using the *C. sporogenes* gene products (YgeX, PbuX, HyuA, YgeW, YgfK, SsnA, YgeY, and XdhAC) and mapped the percent amino acid identities along with uric acid consumption data from the screens onto a phylogenetic tree (**Figure 5A**). Due to incomplete representation of assembled genomes, this analysis included 187 of the 243 strains we tested (**Table S9**). We found that the phylogenetic distribution of uric acid metabolic genes is broad, occurring within 4 phyla, 19 families, and 21 genera. Notably, most organisms that consumed >50% uric acid also contained at least 4/7 uric acid metabolic genes. We also found that most strains that did not contain uric acid metabolic genes also did not consume uric acid. The presence of uric acid metabolic genes also explained differences in uric acid consumption between phylogenetically related bacteria from **Figure 1G** (**Table S10**). When inspecting the genomic context for uric acid metabolic genes, we found that these uric acid metabolic genes mapped to discrete gene clusters across broad taxonomic lineages (**Figure 5B**).

**Figure 5.**
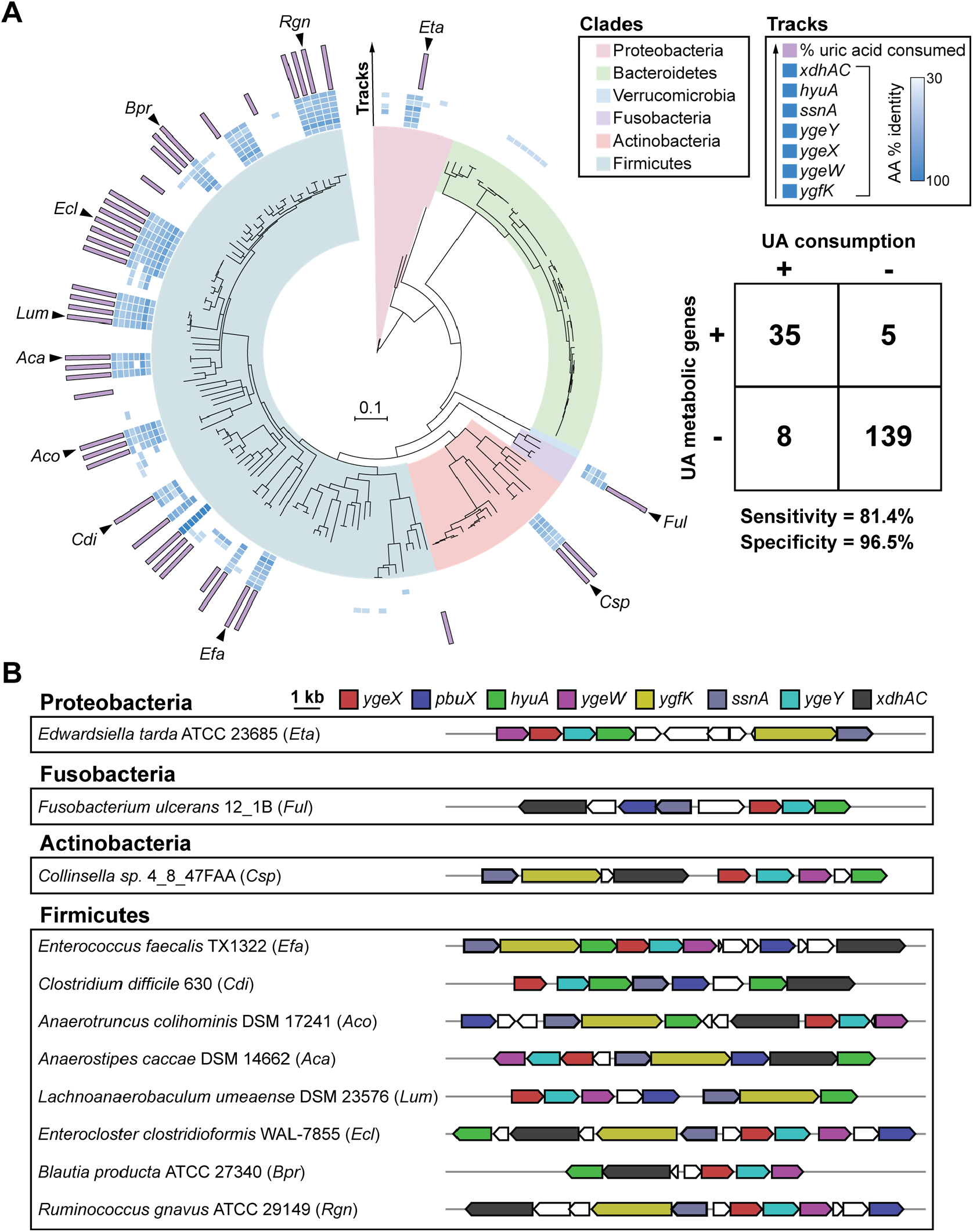
Uric acid gene cluster is conserved across uric acid consuming gut bacteria. A) RpoB phylogenetic tree for strains screened for uric acid metabolism in this study. Only those strains with assembled genomes are included (n = 187). Clades are colored by phylum. Inner blue shaded tracks represent the % amino acid identity of protein homologs identified from BLASTp searches of the relevant genomes using *C. sporogenes* proteins as queries. The outer most track represents the % uric acid consumed by each strain. Uric acid consumption values are only shown for strains with ≥ 50% uric acid consumption. Table shows number of bacteria positive or negative for genes (cut-off ≥ 4 of 7 genes) vs. positive or negative for uric acid consumption (cut-off ≥ 50% uric acid consumption). B) Genomic context of uric acid metabolic genes from representative uric acid consuming strains corresponding to black arrows in Figure 5A.

Within our library, eight strains degraded uric acid but did not harbor the uric acid metabolic genes. Two of these strains included previously studied obligate uric acid degrading bacteria (*Clostridium cylindrosporum* and *Gottschalkia purinilytica*). These organisms are known to convert uric acid to acetate, but our data suggest they use a different set of genes, the identify of which remain unknown. The remaining 6 strains (*Catenibacterium mitsuokai* DSM 15897, *Blautia sp*. KLE 1732, *Dorea formicigenerans* ATCC 27755, *Eubacterium rectale* ATCC 33656, *Coprococcus sp*. HPP0048, and *Veillonella sp*. HPA0037) were among the bacteria we identified as being xanthine producers (**Figure S2** and **Figure 2D**). These organisms likely use xanthine dehydrogenase to convert uric acid to xanthine.

### Escherichia coli converts uric acid to acetate anaerobically

We found that the *E. coli* genome harbors a gene cluster containing most of the uric acid inducible genes identified in our study. *E. coli* has previously been demonstrated to consume uric acid under anaerobic conditions in a mechanism requiring formate and involving several genes including *aegA, ygfT*, and *ygeV* (Iwadate and Kato, 2019). Of note, the *ygfT* and *ygeV* genes map to the same gene locus as the *ygeW, ygeX, ygeY, hyuA, ygfK, ssnA* genes uncovered in our study (**Figure 6A**). However, the role of these latter genes in uric acid metabolism by *E. coli* has not been formally studied.

**Figure 6.**
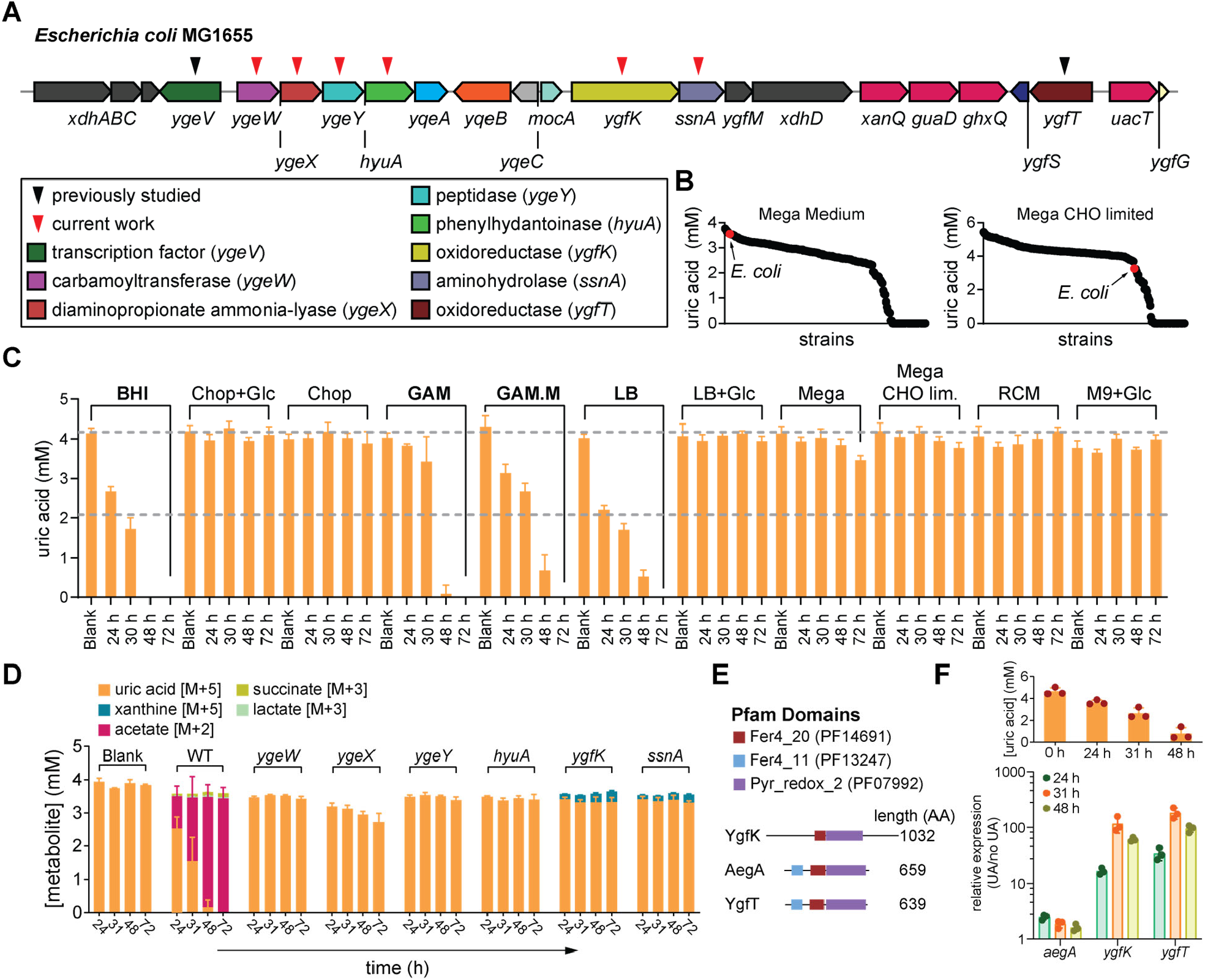
Nutrient dependence of *E. coli* uric acid metabolism and role of genes in conversion of uric acid to acetate. A) Genomic context for uric acid metabolic genes in *E. coli*. Genes indicated by black triangles were previously found to be involved in uric acid consumption. Genes characterized for uric acid metabolism in the current study are indicated by red triangles. B) Results from uric acid metabolism screens under carbohydrate (CHO) replete (left) or CHO limited (right) conditions. Each dot represents an individual strain. Strains are ordered by amount of uric acid remaining and *E. coli* is indicated by a red dot. C) Uric acid metabolism by *E. coli* under different nutrient conditions. *E. coli* was cultured in different media and uric acid levels at the indicated times were quantified by LC-MS. Bold labels indicate conditions where > 50% of uric acid was consumed. BHI, brain-heart infusion broth; Chop, chopped meat medium; Glc, glucose; GAM, Gifu anaerobic medium; GAM.M, modified Gifu anaerobic medium; LB, Luria-Burtani broth; Mega, mega medium; Mega CHO lim., carbohydrate limited mega medium; RCM, reinforced Clostridial medium; M9, M9 defined medium. D) Stable isotope tracing in wild-type and mutant *E. coli* strains. Strains were cultured in modified Gifu anaerobic medium containing either labeled or unlabeled uric acid. Labeled substrates and products were quantified by LC-MS. To account for natural isotope abundances, labeled metabolites detected in unlabeled cultures were subtracted from labeled metabolites detected in labeled cultures. E) Pfam domains for YgfK and two gene products (AegA and YgfT) previously shown to be involved in uric acid metabolism by *E. coli*. F) Relative expression of *ygfK, aegA*, and *ygfT* in uric acid supplemented vs. non-supplemented conditions. Relative expression was determined by RT-qPCR using *rpoH* as a housekeeping gene. Uric acid remaining in the medium is shown in the upper panel. For B, data in the two panels represent the results from a single experiment per condition. For C, D, and F, data represent the means ± standard deviations of n = 3 biological replicates.

Despite containing the genes for uric acid metabolism, under the conditions of our initial screen, *Escherichia coli* was not identified as a uric acid consuming bacterium (**Figure 6B**). To test whether *E. coli* consumes uric acid under anaerobic conditions, we cultured the MG1655 strain in 11 different anaerobic media supplemented with uric acid and monitored uric acid over time by LC-MS. Consistent with results from our initial screen, uric acid was not substantially consumed in mega media (**Figure 6C**). However, we identified 4 different media which supported nearly complete consumption of uric acid (**Figure 6C** and **Table S11**). Comparison of the media formulations suggested that glucose might suppress uric acid metabolism, however this phenotype appears more complex as some media containing glucose (e.g., GAM and BHI) still supported uric acid consumption. Regardless, our findings suggest that *E. coli* does metabolize uric acid and that nutrient availability dramatically influences this phenotype.

Next, we asked whether the uric acid inducible genes identified in our study are required for uric acid metabolism by *E. coli*. To address this, we created markerless deletion mutants in *E. coli* MG1655. We then used stable isotope tracing to quantify uric acid metabolism and conversion to SCFAs by wild-type and mutant *E. coli* MG1655 during growth in modified Gifu anaerobic medium (GAM.M). Under these conditions, the wild-type *E. coli* strain consumed all the uric acid by 48 hours, and culture supernatants accumulated M+2 acetate (**Figure 6D** and **Table S12**). By comparison, the mutant strains were partially blocked in uric acid metabolism and none of the cultures accumulated M+2 acetate (**Figure 6D** and **Table S12**). These findings provide evidence that under certain nutrient conditions, *E. coli* degrades uric acid to acetate, in a pathway that involves *ygeW, ygeX, ygeY, hyuA, ygfK*, and *ssnA*.

Previous studies identified *aegA* and *ygfT* as genes involved in formate-dependent uric acid metabolism in *E. coli* (Iwadate and Kato, 2019). These two genes encode putative oxidoreductases that harbor iron sulfur cluster binding domains and a pyridine-dependent oxidoreductase domain (**Figure 6E**). AegA and YgfT have been proposed to accept electrons from formate dehydrogenase and transfer them to NADP+ or directly to uric acid (Iwadate and Kato, 2019). We found that YgfK also shares two of the three domains present in AegA and YgfT (Fer4_20 and Pyr_redox_2), suggesting that these three enzymes might perform analogous reactions under different nutrient conditions. Given that the conditions of uric acid metabolism in our assay are very different from previous work, we sought to compare the expression of these genes in response to uric acid availability. We cultured *E. coli* MG1655 in medium with or without uric acid and quantified by RT-qPCR the relative expression of these three genes while uric acid was being consumed (**Figure 6F**). The expression of *aegA* was modestly induced under these conditions, however both *ygfK* and *ygfT* were highly induced (**Figure 6F**). These results suggest that both *ygfT* and *ygfK* are likely involved in uric acid metabolism under these conditions, whereas *aegA* is not.

### Gut bacteria protect against hyperuricemia in uricase deficient mice

Next, we sought to address whether uric acid metabolism in the gut might influence levels in the host using a mouse model. Unlike humans, mice have a functional uricase enzyme (also known as urate oxidase (Uox)) and consequently wild-type mice have lower levels of plasma uric acid compared to humans (Lu et al., 2019). To investigate the role of the microbiome in hyperuricemia, we obtained gene edited mice carrying a targeted mutation in the urate oxidase gene (**Figure 7A**) (Wu et al., 1994). These Uox mice have hyperuricemia, accumulate uric acid crystals in the kidney, and suffer from early lethality (Wu et al., 1994). To overcome perinatal lethality, we provided allopurinol (a xanthine oxidase inhibitor) in the drinking water during breeding and after weaning (**Figure 7A**) (Wu et al., 1994).

**Figure 7.**
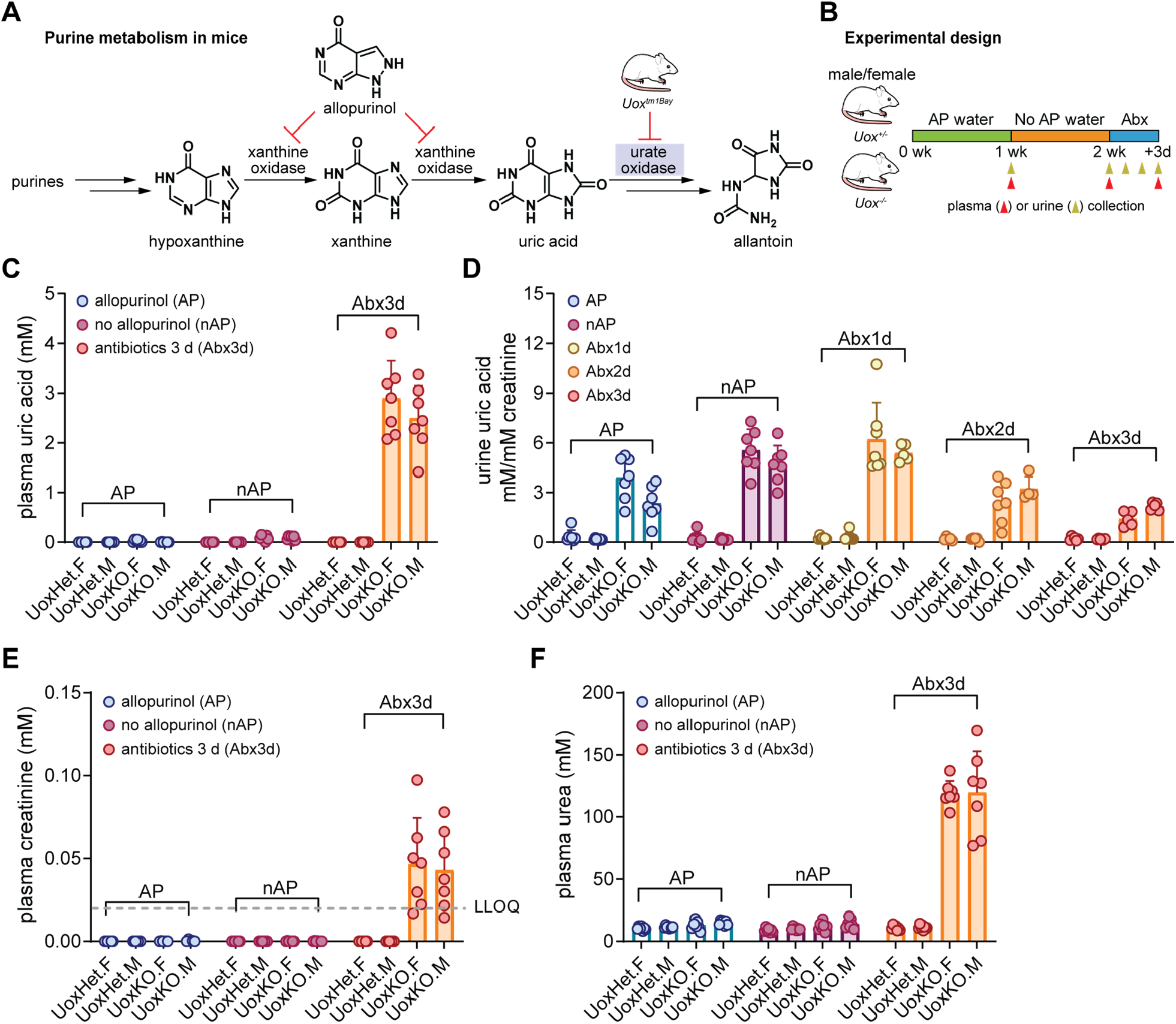
Gut bacteria compensate for loss of uricase in mice. A) Overview of purine metabolism in mice. Mice have a functional urate oxidase (uricase) enzyme. The *Uox*^*tm1Bay*^ mouse line carries a disruption in the urate oxidase gene. Allopurinol, an inhibitor of xanthine oxidase is used to limit hyperuricemia in this mouse model. B) Overview of experimental design. Male (M) or female (F) *Uox*^*tm1Bay*^ (Uox) knockout (KO) or their heterozygous (Het) littermate controls are used. Mice are maintained on allopurinol (AP) water for one week, then given regular water for 1 week, and finally antibiotics were administered in the drinking water for 3 days. Timing of urine and plasma collection are indicated with gold (urine) or red (plasma) arrows. C) Plasma uric acid levels in male and female Uox KO or Het mice. AP, allopurinol; nAP, no allopurinol; Abx, antibiotics. D) Urine uric acid levels normalized to creatinine in male and female Uox KO or Het mice. E) Plasma creatinine levels in male and female Uox KO or Het mice. LLOQ indicates values that are below the lower limit of quantitation. F) Plasma urea levels in male and female Uox KO or Het mice. For C-F, data represent means ± standard deviations from n = 7-8 mice per group.

To test whether the gut microbiota can control uric acid levels in the host, we treated male and female Uox mice and their heterozygous littermate controls with an antibiotic cocktail and measured serum and urine uric acid concentrations. We maintained mice on allopurinol in the drinking water for one week, then removed allopurinol for another week, and finally administered antibiotics in drinking water for 3 subsequent days (**Figure 7B**). After cessation of allopurinol treatment, serum and urine uric acid concentrations in Uox mice increased modestly (**Figure 7C-D**). Unexpectedly, after a three-day course of antibiotics, KO mice but not their heterozygous littermate controls, became ill and developed severe hyperuricemia (**Figure 7C** and **Tables S13-S14**). In the three days after antibiotic administration, urine uric acid excretion progressively decreased, suggesting that kidney function was impaired (**Figure 7D**). Indeed, Uox mice showed evidence of acute kidney injury with markedly elevated concentrations of plasma creatinine and urea (**Figure 7E-F** and **Tables S13-S14**). These findings demonstrate that the gut microbiota compensate for loss of uricase in the host.

### Antibiotics targeting anaerobic gut microbes increase risk for Gout in humans

Given our findings that antibiotic treatment induced severe hyperuricemia in Uox KO mice, we next asked whether antibiotic treatment might be a risk factor for gout in humans. To address this, we compared two commonly prescribed oral antibiotics: i) Clindamycin which is known to target anaerobic microbes, and ii) trimethorprim/sulfamethoxazole (Bactrim) which targets aerobic microbes. We hypothesized that clindamycin might uniquely disrupt uric acid degrading gut bacteria because they are anaerobic microbes. We conducted a retrospective cohort study using electronic health records collected from the Stanford Health Care system between 2015 and 2019. We examined all adult patients (regardless of gout history) and compared rates of new or subsequent gout diagnosis in the years following treatment with ≥ 5 days of oral Clindamycin (N = 7,565) vs. ≥ 5 days of oral Bactrim (N = 23,504). The two groups were similar in age (53.1 vs. 53.4 years), sex distribution (59.6% vs. 59.1% female), and co-morbid illness (average comorbidity score 3.2 vs. 3.5) (**Table S15**). In the unmatched cohort, patients treated with Clindamycin had a higher risk for developing a diagnosis of gout compared to patients treated with Bactrim (Hazard Ratio, 1.18, 95% CI, 1.04-1.34, *P* = 0.00913) (**Figure 8A**). After propensity score matching (N = 6,573 for Clindamycin or Bactrim), the risk for a gout diagnosis increased for patients treated with Clindamycin compared to those treated with Bactrim (Hazard Ratio, 1.3, 95% CI, 1.1-1.54, *P* = 0.00262) (**Figure 8B**). Together, our findings suggest that disruption of the anaerobic gut microbiota by antibiotics increases the risk of developing gout in human patients.

**Figure 8.**
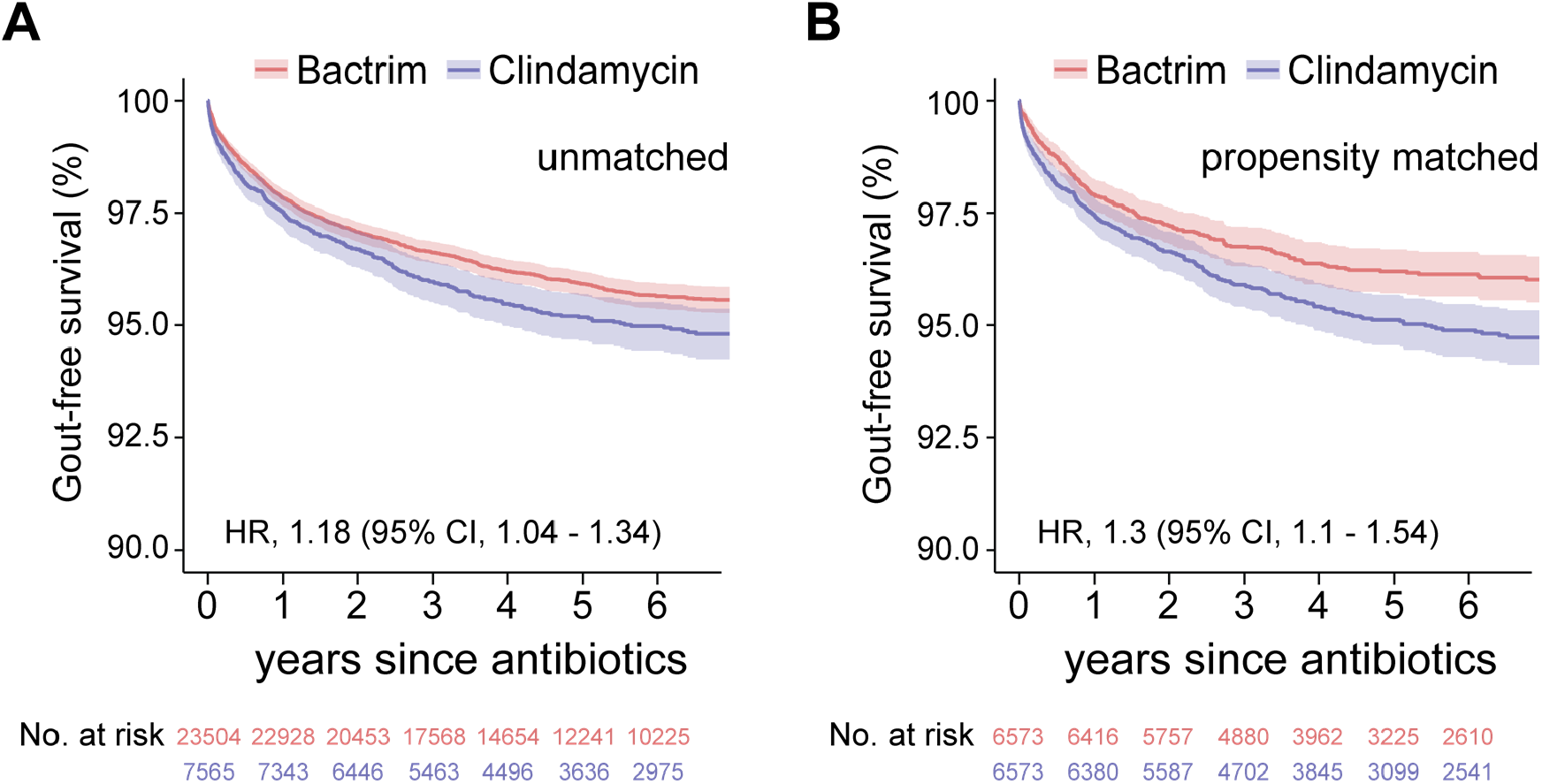
Broad spectrum antibiotics with anaerobic coverage increase risk for gout. Bactrim is a commonly prescribed antibiotic with low coverage against anaerobes, whereas Clindamycin has high anaerobic coverage. A) Kaplan-Meier survival curves for unmatched patients treated with oral Bactrim or Clindamycin (≥ 5 day course) with a diagnosis of gout as the end-point. B) Kaplan-Meier survival curves for propensity score matched patients treated with Bactrim or Clindamycin (≥ 5 day course) with a diagnosis of gout as the end-point.

## Discussion

Uric acid is one of the most abundant nitrogenous compounds on the planet, being a key intermediate in purine metabolism (Barsoum and El-Khatib, 2017). While evidence of aerobic uric acid metabolism can be found across all domains of life, anaerobic uric acid metabolism has been studied in only a handful of bacteria isolated from soil or marine sediments (Vogels and Van der Drift, 1976). Here we find that anaerobic uric acid metabolism is widespread among members of the human gut microbiome, occurring in ∼1/5 of bacteria from 4 of 6 major phyla. In contrast to aerobic pathways that rely on oxygen-dependent uricase to initiate uric acid metabolism, we find that anaerobic pathways break down uric acid through action of uncharacterized ammonia lyase, peptidase, carbamoyl transferase, and oxidoreductase enzymes. The genes encoding these enzymatic functions map to a conserved gene cluster that is broadly distributed across distantly related bacterial taxa and are required for anaerobic uric acid metabolism to lactate and SCFAs. Intriguingly, previously characterized uric acid degrading Clostridia (e.g., *G. purinilytica, G. acidiurici*, and *C. cylindrosporum*) do not encode these genes (Hartwich et al., 2012; Poehlein et al., 2015a; Poehlein et al., 2015b), suggesting that distinct pathways for anaerobic uric acid metabolism evolved independently among bacteria. However, the uric acid genes identified in our study are highly predictive of uric acid metabolism activity in gut bacteria, indicating this gene cluster encodes a predominant pathway for anaerobic uric acid metabolism in the gut.

Uric acid is a major nitrogenous waste product in birds, insects, reptiles, and land snails. Evidence suggests that uric acid is also held as a metabolic reserve in birds and termites (Potrikus and Breznak, 1981; Singer, 2003), and that intestinal bacteria play a role in nitrogen recycling from uric acid when the host subsists on a nitrogen-poor diet. However, in both systems, the bacteria and genes responsible for uric acid breakdown (uricolysis) are unknown. We analyzed metagenome assembled genomes (MAGs) from chicken feces (Gilroy et al., 2020) and the termite hindgut (Herve et al., 2020) and found that many organisms harbor the uric acid gene cluster (subset shown in **Figure 9**). Thus, we find that the uric acid gene cluster is distributed broadly across different ecosystems on the planet and may play an important role in global nitrogen cycling.

**Figure 9.**
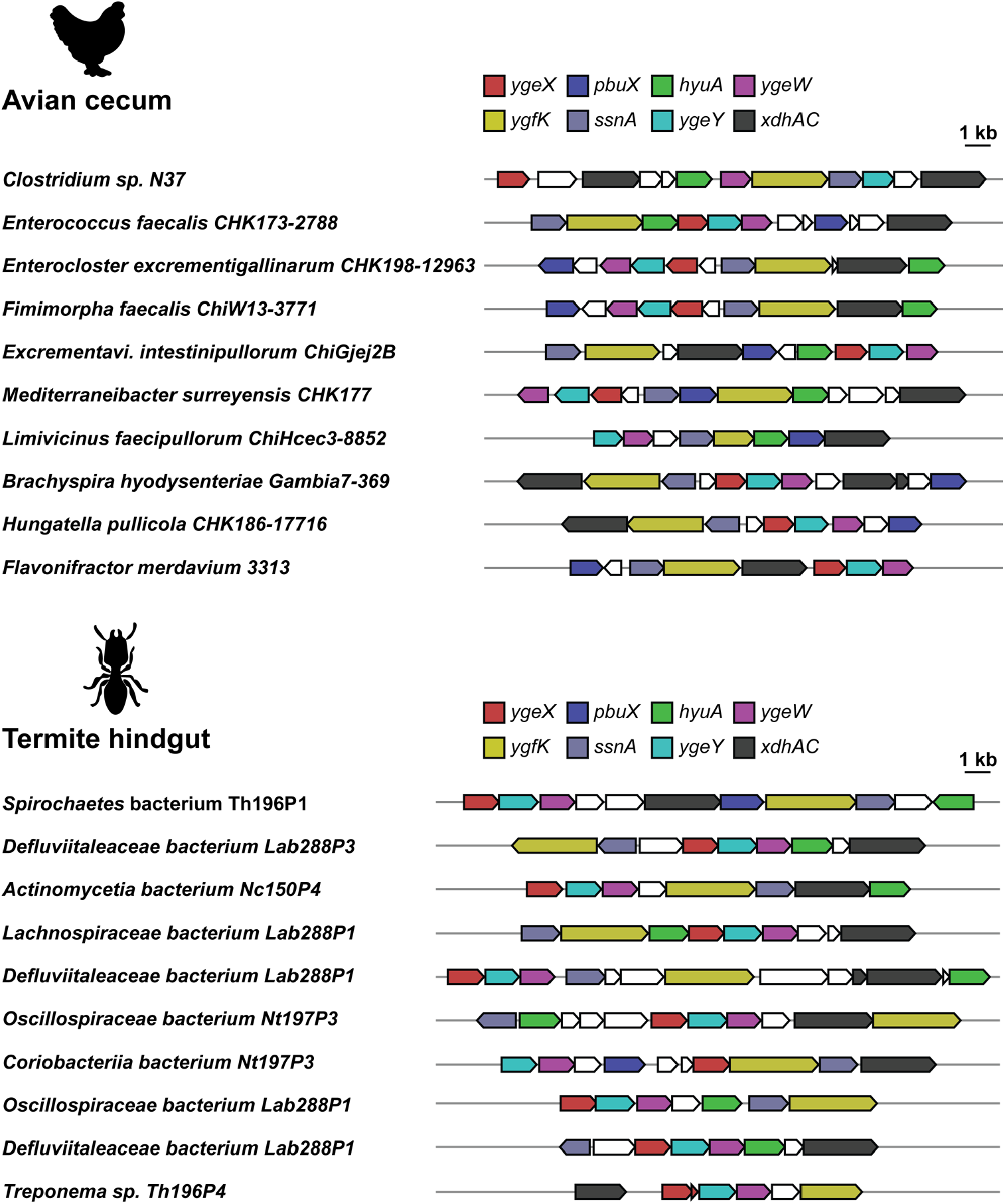
Uric acid gene cluster is present in metagenome assembled genomes from the gut of uricotelic organisms. Both birds and termites are uricotelic, excreting uric acid as their primary nitrogenous waste product in their gut. The uric acid gene cluster identified in this study was used to search metagenome-assembled genomes from the chicken cecum (NCBI BioProject PRJNA543206) or the termite hindgut (NCBI BioProject PRJNA560329). The genomic context for uric acid metabolic genes is shown for representative taxa from these two metagenomic studies.

In most mammals purines are degraded via uricase to freely soluble allantoin which is excreted in the urine. However, uricase was inactivated early in hominid evolution predisposing humans to hyperuricemia and gout. It has long been known that uric acid is eliminated in the gut and subject to uricolysis by gut bacteria (Geren et al., 1950; Lucke, 1931a, b; Sorensen, 1965; Thannhauser and Dorfmüller, 1918). However, it is not known whether microbial uric acid metabolism in the gut contributes to uric acid homeostasis in the host. By modeling uricase deficiency in humans with uricase-deficient mice, we found that disruption of gut microbiota with antibiotics causes profound hyperuricemia. After just 3 days of antibiotic treatment these uricase-deficient mice become ill, developing acute kidney injury likely due to accumulation of uric acid crystals in the kidney (Wu et al., 1994). We also find that patients given clindamycin – an antibiotic known to target anaerobic gut bacteria – are at higher risk for developing gout relative to those treated with antibiotics that have limited anaerobic coverage. The increased risk is intermediate in magnitude (hazard ratio 1.3, Clindamycin vs. Bactrim) which may explain why this connection has not been previously identified in the clinic. Together our findings support a model whereby the gut microbiota, through the capacity for uric acid metabolism, compensates for the loss of uricase activity in humans.

To address whether disruption of gut bacteria influences fecal uric acid levels, we re-analyzed metabolomics data from a recent human study which characterized microbiota recovery after depletion with antibiotics and polyethylene glycol (PEG) (Tanes et al., 2021). In this study, microbiota depletion resulted in dramatically elevated fecal levels of uric acid (**Figure S4**). Fecal uric acid levels rapidly returned to baseline in subjects fed a vegan or omnivore diet, but those fed a fiber-free synthetic diet showed a protracted recovery and several study subjects showed persistent elevations of fecal uric acid throughout the recovery phase (**Figure S4**). The authors of this study demonstrated a delay in recovery of members of the Firmicutes in the fiber-free diet (Tanes et al., 2021) which is consistent with our observation that Firmicutes are dominant uric acid degrading bacteria in the gut. This suggests that a lack of dietary fiber following microbiome perturbation imparts a sustained dysregulation of uric acid metabolism in the gut.

Our study provides new insights into the role of the gut microbiota in hyperuricemia and gout, two common disorders affecting the US population. There are two important implications of these results: First, antibiotic therapies that might disrupt the gut microbiota should be carefully considered in patients predisposed to gout. If antibiotics are administered to these patients, a low fiber diet may cause a protracted return to normal uric acid metabolism in the gut. Second, increasing uric acid metabolism in the gut represents an important new therapeutic approach to treating patients with hyperuricemia. Along these lines, recent studies have shown that oral (non-absorbable) enzyme therapy with recombinant uricase reduces plasma uric acid concentration in uricase-deficient mice (Pierzynowska et al., 2020), and decreases plasma concentration in healthy volunteers (Terkeltaub et al., 2022). We envision that live biotherapeutic products consisting of uric acid consuming bacteria could also be an important new therapeutic modality to treat hyperuricemia and gout.

## Methods

### Reagents used in this study

All chemicals and reagents used in this study were of the highest possible purity and are listed in **Table S16**. Uniformly labeled [^13^C_5_] uric acid was synthesized by Acanthus Research Inc. (Mississauga, Ontario, Canada). The analytical summary including ^1^H-NMR, ^13^C-NMR, high resolution mass spectrometry, are provided in the Supplementary material. It is available for purchase from Acanthus Research Inc. as catalog # U-10826-01. Due to solubility issues, uric acid or its ^13^C_5_ isotope was made fresh as follows: stock solutions of uric acid were prepared at 125 mM in 1 N NaOH (for screening) or 120 mM in 0.4 N NaOH (for all other experiments), sterile filtered and then diluted appropriately in anaerobic media for uric acid consumption assays.

### Bacterial strains and culture conditions

Bacterial strains used in this study were obtained from culture collections as listed in **Table S17** and medium formulations are provided in **Table S18a-e**. All bacteria were cultured at 37 °C in a Coy type B anaerobic chamber using a gas mix containing 5% hydrogen, 10% carbon dioxide, and 85% nitrogen. An anaerobic gas infuser was used to maintain hydrogen levels of 3.3%. All media and plasticware were pre-reduced in the anaerobic chamber for at least 24 hours before use. *E. coli* for genetic manipulation was cultured under aerobic conditions using LB broth and LB plates, with temperature and antibiotic selection varying depending on the manipulation being done. *E. coli* uric acid consumption was performed under anaerobic conditions using pre-reduced media.

### Culture conditions for uric acid consumption assays

Bacterial strains used in the library, along with their culture media, are listed in **Tables S1-S2**. All strains were stored at -80 °C as anaerobically prepared 20% glycerol stocks, sealed to ensure anoxic conditions for long term storage. All bacteria were cultured in 96-deep well plates anaerobically unless otherwise indicated. For bacteria library screening, glycerol stocks were first inoculated in rich media without uric acid at 37 °C for 24 h. Cultures were then diluted (10-fold) into medium containing uric acid (5 mM) and continued to incubate for 48 h. The culture supernatant was harvested by centrifugation at 5,000 *g* for 25 min, 4 °C. Aliquots of supernatants were transferred to 96-well microtiter plates, tightly sealed, and stored at -80 °C prior to sample preparation for LC-MS analysis. For other *in vitro* culture assays, bacteria strains were first streaked on GAM or RCM plates, and individual well isolated colonies were picked to inoculate in medium broth. *E. coli* was cultured in modified GAM broth unless otherwise indicated. Other strains were cultured in GAM broth. Individual colonies were picked and inoculated in rich media with 2 mM uric acid for 16-18 h and then diluted 50-fold in media with 4 mM uric acid. At designated time points, aliquots of cultures were harvested by centrifugation (5,000 *g*, room temperature, 5 min). Supernatants were collected and aliquoted into two plates, one that was used for SCFA measurement, and the other that was mixed with NH_4_OH (final 10 mM) and used for uric acid measurement. Both plates were stored at -80 °C until analysis by LC-MS.

### Stable isotope tracing with ^13^C_5_ labeled uric acid

For stable isotope tracing, strains were first cultured in rich medium with unlabeled uric acid (2 mM) for 16-18 h before being diluted into medium supplemented with either unlabeled uric acid (4 mM) or uniformly ^13^C_5_ labeled uric acid (4 mM). At designated times, aliquots were harvested as described above. When cultured without labeled uric acid, isotopes (e.g., M+2 acetate and M+2 butyrate) were detectable in some of the cultures. We found that this reflected natural abundance of these isotopes resulting from the large amount of short chain fatty acids produced from nutrients (amino acids and carbohydrates) present in the rich medium. Therefore, to correct for natural isotope abundance, we cultured organisms with labeled and unlabeled uric acid. After quantifying SCFA isotopes, we subtracted concentrations of isotopes in cultures with unlabeled uric acid from those cultures with labeled uric acid.

### Sample preparation for analysis of uric acid, xanthine, and creatinine by liquid chromatography – mass spectrometry (LC-MS)

Uric acid or xanthine was made fresh at 120 mM in 0.4 N NaOH each time. Creatinine was dissolved at 500 mM in LC-MS grade water and stored at -20 °C. The stock solutions were diluted in 10 mM aqueous NH_4_OH to serve as a calibration standard for LC-MS assays. To account for matrix effects, the same portion of medium or charcoal treated human plasma was added to the standard curves for *in vitro* samples or mouse plasma samples, respectively. Standard curves were treated the same as samples during LC-MS preparation.

Culture supernatants, mouse plasma, mouse urine and calibrants were first mixed with internal standard (ISTD) and 10 mM NH_4_OH, and then filtered by AcroPrep Omega 3K MWCO filter plates (Pall Corporation, 8033) at 3,000 *g* for 30 min at room temperature. The flowthrough was collected and diluted in NH_4_OH (final 4 mM) before subjecting to LC-MS analysis.

### Sample preparation for analysis of short chain fatty acids by LC-MS. Culture

supernatants were first mixed with internal standard (ISTD) in a V-bottom, polypropylene 96-well plate, and then extracted by mixing with extraction solution (75% acetonitrile/25% methanol) at 1:3 ratio. The plate was covered with a lid and centrifuged at 5,000 *g* for 15 min at 4 °C. Supernatant was collected for 3-nitrophenylhydrazine derivatization before subjecting to LC-MS analysis.

### 3-Nitrophenylhydrazine (NPH) derivatization protocol

This derivatization method targets compounds containing a free carboxylic acid. Extracted samples were diluted in 50% acetonitrile and then mixed with 3-nitrophenylhydrazine (200 mM in 80% acetonitrile) and *N*-(3-dimethylaminopropyl)-*N*’-ethylcarbodiimide (120 mM in 6% pyridine) at 2:1:1 ratio. The plate was sealed with a plastic sealing mat (Thermo Fisher Scientific cat. # AB-0566) and incubated at 40 °C, 600 rpm in a thermomixer for 60 min to derivatize the carboxylate containing compounds. The reaction mixture was quenched with 0.02% formic acid in 20% acetonitrile/water before LC-MS analysis.

### Quantification of metabolites by liquid chromatography-mass spectrometry (LC-MS)

During the course of this study, two different LC-MS conditions were used (C18 positive underivatized and 3-nitrophenylhydrazine derivatized C18 negative). An overview of the general method is provided here and the specific instrument parameters for the different analytical methods are provided in **Table S19**. Samples were injected via refrigerated autosampler into mobile phase and chromatographically separated by an Agilent 1290 Infinity II UPLC and detected using an Agilent 6545XT Q-TOF equipped with a dual jet stream electrospray ionization source operating under extended dynamic range (EDR 1700 m/z). MS1 spectra were collected in centroid mode, and peak assignments in samples were made based on comparisons of retention times and accurate masses from authentic standards using MassHunter Quantitative Analysis v.10.0 software from Agilent Technologies. Compounds were quantified from calibration curves constructed with authentic standards using isotope-dilution mass spectrometry with appropriate internal standards (**Table S20**).

### RNA purification for RNA-seq experiment

All cultures were grown at 37 °C under anaerobic conditions. *Clostridium sporogenes* ATCC 15579, *Collinsella aerofaciens* ATCC 25986, and *Lacrimispora saccharolyticum* WM1 were streaked from frozen stocks onto blood agar plates. Three individual colonies were selected for each bacterium and were inoculated into separate overnight cultures in Mega medium. The following day, each culture was precultured in Mega medium, and after three hours diluted to an OD of 0.1 into two experimental cultures, one containing standard Mega medium and one with Mega medium containing 5 mM uric acid. Bacteria were allowed to grow until reaching an OD that was commensurate with approximately 50% uric acid degradation as determined by previous experiments. Cell cultures were then combined with two volumes of RNAprotect (Qiagen) in an anaerobic chamber, mixed thoroughly and then allowed to sit for five minutes. Cells were then centrifuged (5,000 *g*, 4 °C, 10 min) and the supernatant was decanted. Cells were then subjected to lysozyme digestion, Proteinase K digestion and mechanical disruption with a mixer mill (RETSCH MM400) at 4 °C, 25/s, for 30 min. RNA was then purified using RNeasy kit (Qiagen), followed by DNase digestion and second RNA purification step using the RNeasy kit (Qiagen). RNA integrity was determined using a bioanalyzer (Agilent) and RNA-seq was performed by the Roy J. Carver Biotechnology Center at the University of Illinois.

### RNA-seq library preparation and data collection

Ribosomal RNA was removed with the Ribo-Zero Bacteria kit (Illumina). The RNA-seq libraries were prepared using a TruSeq Stranded mRNAseq Sample Prep kit with each sample individually ligated with unique adapters (Illumina). The libraries were quantitated by qPCR, pooled, and sequenced on one lane for 101 cycles from one end of the fragments on a NovaSeq 6000 using a NovaSeq SP reagent kit. Fastq files were generated and demultiplexed with the bcl2fastq v2.20 Conversion Software (Illumina) and adaptors were removed from the 3’ end of the reads. Read 1 aligns to the antisense strand and read 2 aligns to the sense strand.

### RNA-seq data analysis

RNA-seq processing was performed using CLC Genomics Workbench (v.21.0.4). Reads were trimmed using a quality score limit of 0.05 and ambiguous nucleotides (n = 2), and automatic read-through adapter trimming was performed. Next, genomes were downloaded from NCBI in GenBank file format (.gbff) for each of the three bacteria (NCBI assembly accession numbers: GCF_010509075.1, *Collinsella aerofaciens* ATCC 25986; GCF_000144625.1, *Lacrimispora saccharolyticum* WM1; GCF_000155085.1, *Clostridium sporogenes* ATCC 15579). Genomes were uploaded into CLC Genomics Workbench and converted to tracks. RNAseq was performed using the genome track as the reference sequence and genes for the gene track. Mapping settings included: Mismatch cost, 2; Insertion cost, 3; Deletion cost, 3; Length fraction, 0.8; Similarity fraction, 0.8; maximum number of hits for a read, 10. Expression settings included: Strand setting, both; Library type setting, bulk; Expression value, TPM. Statistical comparisons were made between organisms cultured with or without uric acid using multi-factorial statistics based on a negative binomial GLM as implemented in CLC Genomics Workbench (v.21.0.4). The expression and CDS tracks were then exported as Excel files and expression and annotation tracks were merged in Excel using the VLOOKUP function based on the chromosome coordinates. For Volcano plots, the -Log_10_ False Discovery Rate (FDR) corrected *P*-value was plotted against the Log_2_ fold-change for cultures grown with vs. without uric acid.

### Gene disruption in Clostridium sporogenes ATCC 15579 using ClosTron

*Plasmid construction and cloning*. For *Clostridium sporogenes* gene disruptions were constructed using the Intron targeting and design tool on the ClosTron website (http://www.clostron.com/clostron2.php) with the Perutka algorithm (Heap et al., 2010). The intron within the pMTL007C-E2 plasmid was targeted to the sites listed in **Table S21** and the targeting sequences were synthesized by IDT. New targeting plasmids were made by digesting the pCsp-316-736s (Dodd et al., 2017) plasmid with BsrGI and HindIII, followed by gel-purification of the plasmid backbone and Gibson assembly with re-targeted intron fragments using the Gibson Assembly Master Mix (NEB). Gibson assemblies were transformed into *E. coli* by electroporation, selected on LB-chloramphenicol (25 µg/mL) plates and sequenced to confirm the correct retargeted sequence. Intron re-targeted plasmids were transformed into *E. coli* s17-1 *λpir* and subsequently conjugated into *C. sporogenes* as described before (Liu et al., 2022). Mutants were verified by PCR and Sanger sequencing using the sequencing primers listed in **Table S21**.

### Gene disruption in Escherichia coli MG1655 using λ-red recombination

Gene disruption in *Escherichia coli* was first constructed by the Red recombination deletion method (Datsenko and Wanner, 2000) in BW25113 strain and then was transferred to MG1655 by P1 transduction (Thomason et al., 2007). The antibiotic resistance cassettes were removed by FLP-mediated excision (Datsenko and Wanner, 2000) before being used in experiments. Mutants were verified by PCR using primers listed in **Table S21**. All strains used in this study are listed in **Table S22**.

### RNA purification for RT-qPCR

All cultures were grown at 37 °C under anaerobic conditions. *Escherichia coli* MG1655 was streaked out from frozen stocks onto GAM plates. Individual colonies were selected and were inoculated into separate cultures in GAM modified medium with or without 2 mM uric acid. Three individual cultures were used for each condition. After 16 hours, the overnight cultures were diluted 50-fold in GAM modified medium with or without 4 mM uric acid and continued to incubate at 37° C. At 24 h, 31 h and 48 h, one aliquot of culture was saved for uric acid LC-MS measurement, and another aliquot of culture was mixed with two volumes of RNAprotect Bacteria Reagent anaerobically to stabilize the RNA. RNA was extracted as described above. The total RNA concentration was measured by Qubit RNA BR Assay Kit. Two micrograms of total RNA were used for ezDNase digestion and then was reverse transcribed to cDNA by SuperScript IV VILO in a 20 µL reaction following manufacturer’s guidelines. Q-PCR was performed using PowerUp SYBR Green Master Mix with 4 replicates and an Applied Biosystems QuantStudio™ 5 real-time PCR instrument (ThermoFisher). *E. coli* gDNA concentration was measured using Qubit dsDNA BR Assay Kit. Primer validation was performed using six serial 10-fold dilutions of *E. coli* gDNA, spanning 2 ng/µL to 0.002 pg/µL per reaction. Primer amplification factor (Ep) and efficiency were calculated by ThermoFisher qPCR Efficiency Calculator. Three housekeeping genes (*dnaK, fliA* and *rpoH*) were tested and *rpoH* was selected as the reference gene because its Ct was consistent regardless of uric acid addition and was most similar to the Ct of target genes. Relative fold change of the target gene was calculated as follows:

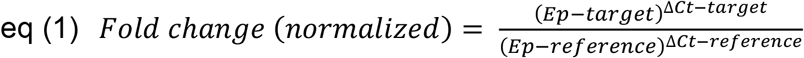

### Uox mice experiment

Uox (B6; 129S7-Uox^tm1Bay/J^) mice were resuscitated from frozen embryos by The Jackson Laboratory (strain # 002223). Animal experiments were performed following a protocol approved by the Stanford University Administrative Panel on Laboratory Animal Care. The mouse strain was maintained by heterozygous female (+/-) x homozygous male (-/-) mating. Mice were kept on standard chow (LabDiet Cat. # 5K67) and allopurinol water (100 mg/L) with access to food and water ad libitum in a facility on a 12-hour light/dark cycle with temperature controlled between 20-22°C and humidity between 40-60%. Genotyping was performed using Terra PCR Direct Genotyping Kit (Takara, 639285) following protocol modified from The Jackson Laboratory. For antibiotic treatment, water was supplemented with 0.5 mg/mL vancomycin, 1 mg/mL neomycin, 1 mg/mL metronidazole and 1 mg/mL colistin. Blood sampling was performed via the facial vein, collecting ∼100 µL of blood into tubes containing concentrated sodium EDTA (final ∼12 mM) as an anticoagulant and allopurinol (final ∼12 µM) to inhibit xanthine oxidase (Watanabe et al., 2014). After centrifugation at 1,500 *g* for 10 min at 10 °C, plasma was transferred to new tubes. Urine was collected by manually expressing urine from individual mice into sterile tubes. All samples were stored at -80 °C. Uric acid and creatinine were measured by LC-MS, and urea was measured by Urea Assay Kit.

### Impact of antibiotic treatment on risk for Gout diagnosis

We conducted a retrospective new user cohort study (Ray, 2003) using electronic health records (EHR) collected during from the Stanford Health Care system between 2015 and 2019. EHR were mapped to the Observational Medical Outcomes Partnership (OMOP) Common Data Model (CDM) version 5.3 such that standardized vocabularies like the RxNORM and International Classification for Diseases (ICD) could be used to define patients and their conditions (**Table S23**). Patients between 18 and 90 years old were included in the cohort if they were prescribed at least 5 days of oral Bactrim and Clindamycin and if they did not receive antibiotics in the preceding 3 months. Primary outcome was gout diagnosis up to 5 years after antibiotic treatment (see **Table S23** for detailed definitions). The study only included deidentified data and qualified for exempt status by the Institutional Review Board of Stanford University.

### Propensity score model

We created a multivariable logistic regression model to calculate the probability of patients receiving Clindamycin or Bactrim. The propensity score (PS) regression model controlled for the following pre-exposure confounders: age, sex, race/ethnicity, Charlson comorbidities (Quan et al., 2005), diagnostic, procedure and medication codes, and number of encounters observed in the 90 days before antibiotics initiation.

The propensity score provides a composite score of the baseline confounders such that when PS is balanced (within a caliper of 0.25) between the Bactrim and Clindamycin arms, their baseline confounders would also become balanced (Austin, 2011). We used high dimensional propensity scores (hdPS) (Schneeweiss et al., 2009). First, hdPS covariates were generated from ICD, CPT and RxNORM codes. A model of the hdPS covariates is fitted using a logistic regression with LASSO regularization that penalizes low weight covariates down to zero weights such that the resulting parsimonious model has equivalent predictive performance without overfitting too many covariates in a high dimensional setting (Franklin et al., 2015; Low et al., 2016). The LASSO hyperparameter is tuned using 5-fold cross-validation and the 1-standard error rule (Hastie et al., 2009). A two-sided *P*-value of 0.05 was considered statistically significant. Analyses were performed using R version 4.05 on the Atropos Health platform (Callahan et al., 2021).

## Data and Code Availability

The RNA-seq data has been uploaded to the NCBI Gene Expression Omnibus under accession number GSE206419. Bacterial genome assemblies analyzed in this study (e.g., for phylogenetic trees and BLASTp searches) are from publicly available sources and relevant accession numbers are provided in **Table S17**. Metagenome assembled genomes analyzed in this study from termite hindgut and chicken ceca are from publicly available sources (chicken cecum, NCBI BioProject PRJNA543206; termite hindgut NCBI BioProject PRJNA560329). Metabolomics data re-analyzed from a study investigating microbiota recovery after depletion (Tanes et al., 2021) was obtained from the Metabolomics Workbench (Study ID ST001519). No custom code was used in this study.

## Acknowledgments

We are grateful to Justin Sonnenburg, Isaac Cann, and Michael Fischbach for valuable discussions. We thank Arianna Burke, Alejandra Dimas, Steven Higginbottom, and Duy Nguyen for assistance with gnotobiotic animal experiments. We thank James Imlay for providing P1 phage. We also thank Alvaro Hernandez and Chris Wright from the Roy J. Carver Biotechnology Center at the University of Illinois for help with RNA-seq data collection. This work was funded in part by National Institutes of Health grants K08-DK110335 (D.D.), R35-GM142873 (D.D.), R01-AT011396 (D.D.), and an OHF-ASN Foundation for Kidney Research Career Development Award (D.D.).

## Author contributions

Y.Liu, J.B.J., and D.D. conceived and designed the project; Y.Liu and J.B.J. conducted and analyzed uric acid screening assays with help from H.C., D.D., and B.H.; Y.Liu performed in vitro uric acid consumption assays and stable isotope tracing with help from D.D.; J.B.J. performed RNA-seq experiments and analyzed data with help from D.D.; J.B.J. created bacterial mutants with help from Y.Liu, K.S., and D.D.; H.C. developed LC-MS assays and analyzed LC-MS data with help from Y.Liu, J.B.J., and D.D.; Y.Liu prepared samples for LC-MS analysis; S.H. and D.D. conducted phylogenetic analyses; Y.Liu performed RT-qPCR experiments; M.E.D. assisted with breeding of Uox deficient mice; Y.Liu conducted mouse experiments with help from M.E.D., K.S., and D.D.; A.C.P., C.G., D.D., S.G., and Y.S.L. conceived and designed Gout cohort analysis; Y.S.L. conducted cohort analysis, advised by S.G.; Y.Liu and D.D. wrote the manuscript with input from all co-authors; Y.Liu and D.D. prepared all figures with help from S.H., S.G., and Y.S.L.

## Declarations of interest

D.D. is a co-founder of Federation Bio.

