## Supplementary Material for "A widely distributed gene cluster compensates for uricase loss in hominids"

[1]Department of Pathology, Stanford University School of Medicine; [2]Atropos Health, Palo Alto; [3]Department of Cellular and Molecular Biology, University of California, Berkeley; [4]Department of Microbiology and Immunology, Stanford University School of Medicine; [5]Division of Nephrology, Department of Medicine, Stanford University School of Medicine; [6]Department of Urology, Stanford University School of Medicine; [7]Veterans Affairs Palo Alto Health Care System

[†] These authors contributed equally

Supplementary Figures

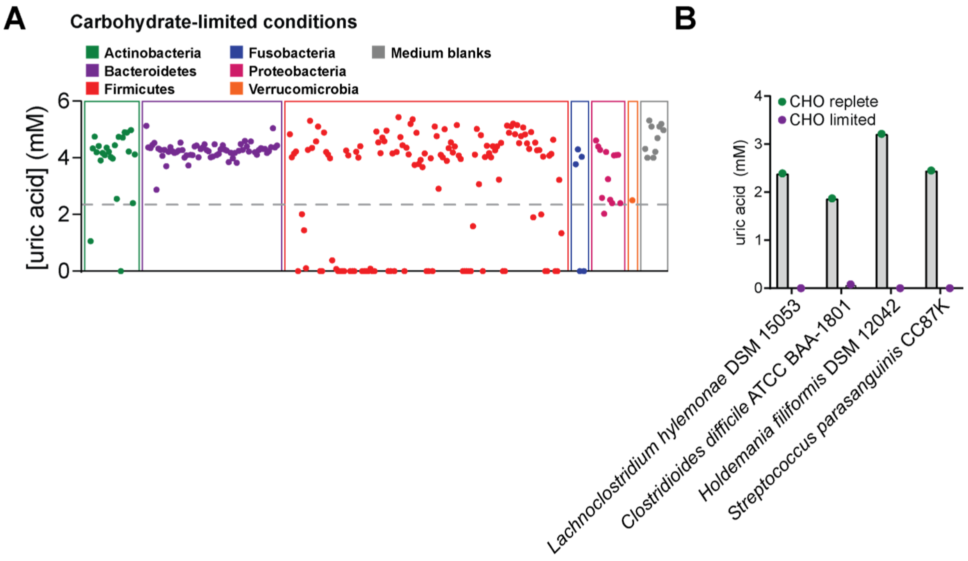

**Figure S1. Carbohydrate availability influences uric acid metabolism.** A) Bacteria were cultured for 48 h in rich medium with limited carbohydrates (Table S2) and uric acid was quantified by LC-MS. Each dot represents a single bacterial strain and organisms are grouped by phylum. B) Bacteria that degrade uric acid in carbohydrate limited medium, but not in carbohydrate supplemented medium. Strains are shown that consumed <50% uric acid in carbohydrate supplemented medium and consumed >50% uric acid when carbohydrates were limited. For A and B, data represent the results from a single experiment.

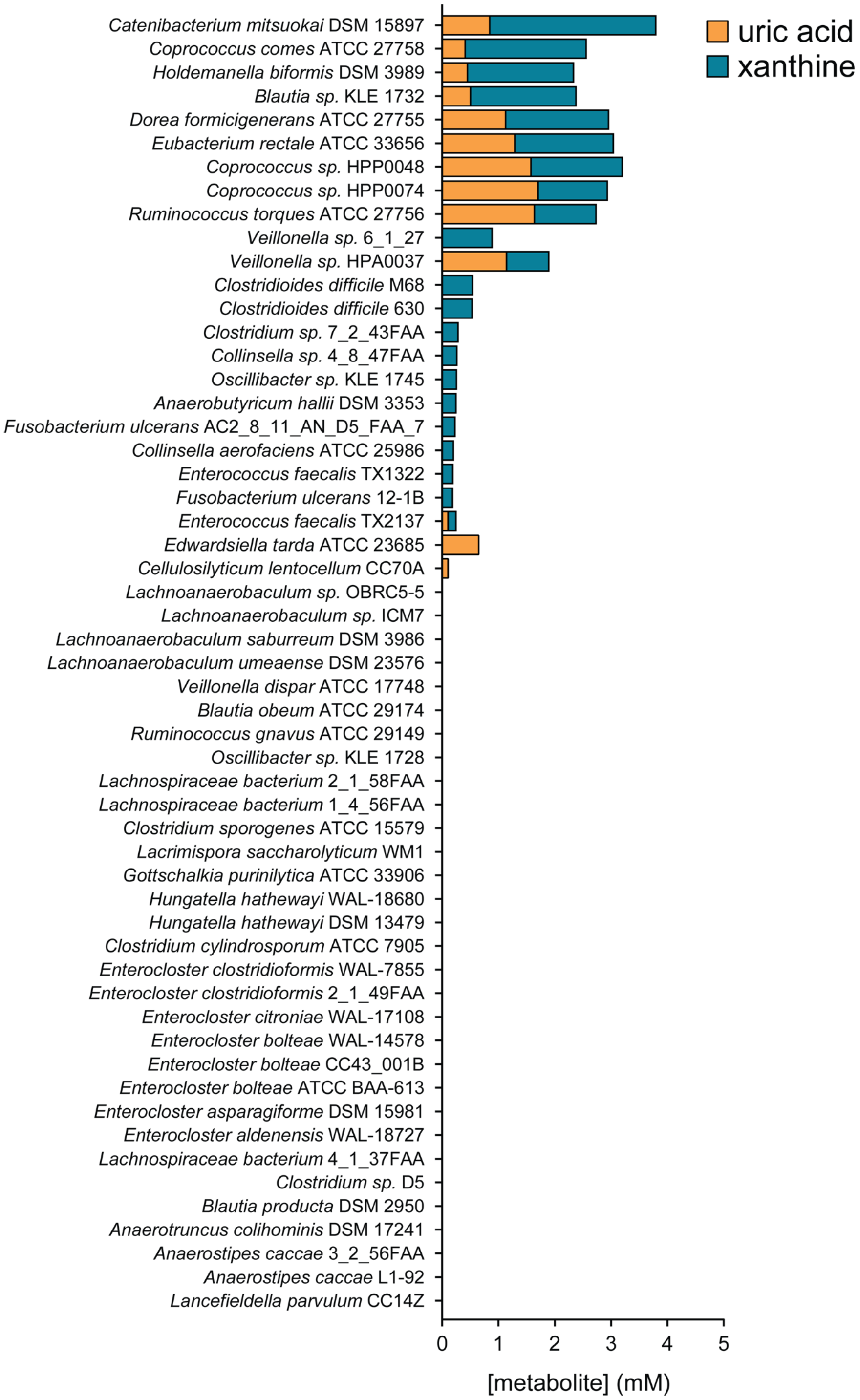

**Figure S2. A subset of gut bacteria accumulate xanthine when consuming uric acid.** Xanthine and uric acid were quantified in the supernatant of both screening experiments after 48 h by LC-MS. Only those strains that consumed ≥ 50% uric acid are shown. Data represent the results from a single experiment.

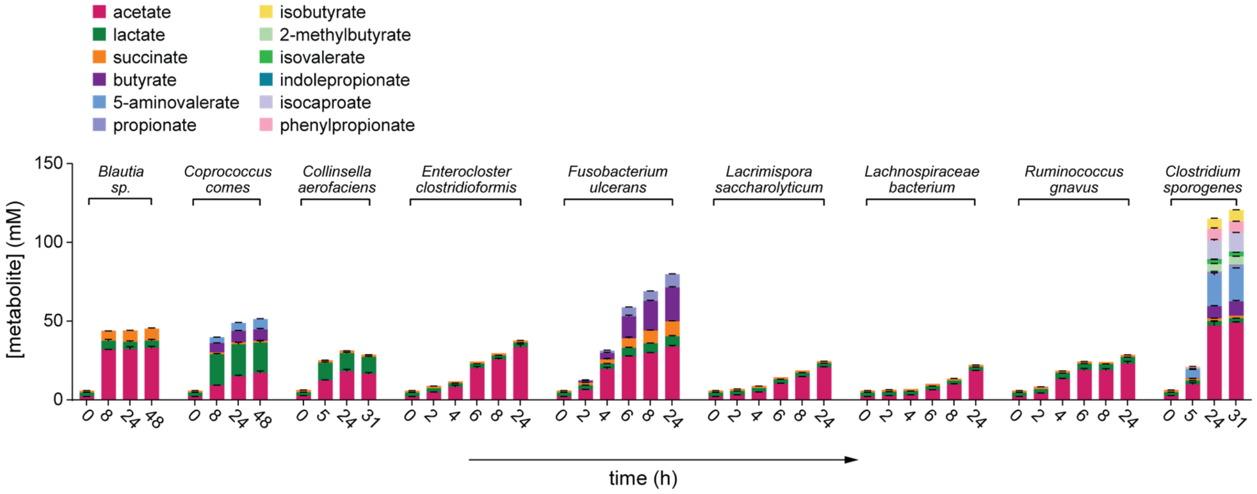

**Figure S3. Short chain fatty acid production by gut bacteria.** Bacteria were cultured in rich medium supplemented with unlabeled uric acid and short chain fatty acids were quantified by LC-MS at the indicated timepoints. Data represent the means ± standard deviations of n = 3 biological replicates.

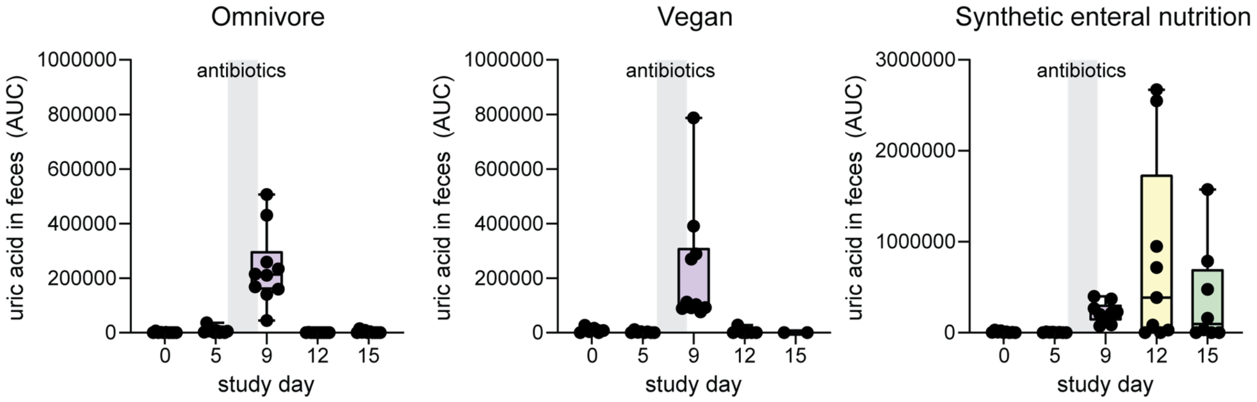

**Figure S4. Microbiota depletion in human subjects increases uric acid levels in the feces.** Data were re-analyzed from a previously published study exploring the role of diet in microbiome recovery after disruption with antibiotics and polyethylene glycol (PEG) (Metabolomics Workbench Study ID: ST001519). Fecal uric acid levels for each study subject are plotted on each day of the study for participants in either the omnivore, vegan, or fiber-free synthetic enteral nutrition groups. Grey region indicates time in which antibiotics and PEG were administered to study participants. AUC, area under the curve.

### Analytical Summary

| | | |
|---|---|---|
| Product Number: | U-10826-01 | 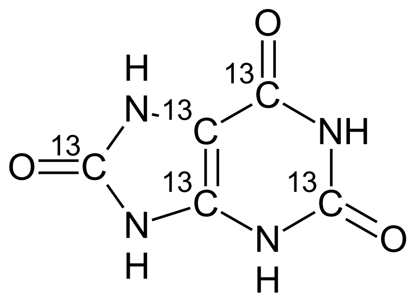 |
| Product Name: | Uric Acid-$^{13}C_5$ | |
| Lot Number: | URAL-1790-P103-72D | |
| Molecular Formula: | $^{13}C_5H_4N_4O_3$ | |
| Molecular Weight: | 173.07 g $mol^{-1}$ | |
| Chemical Name: | 7,9-Dihydropurin-2,6,8-trione-$^{13}C_5$ | |

$^1$H-NMR Spectrum

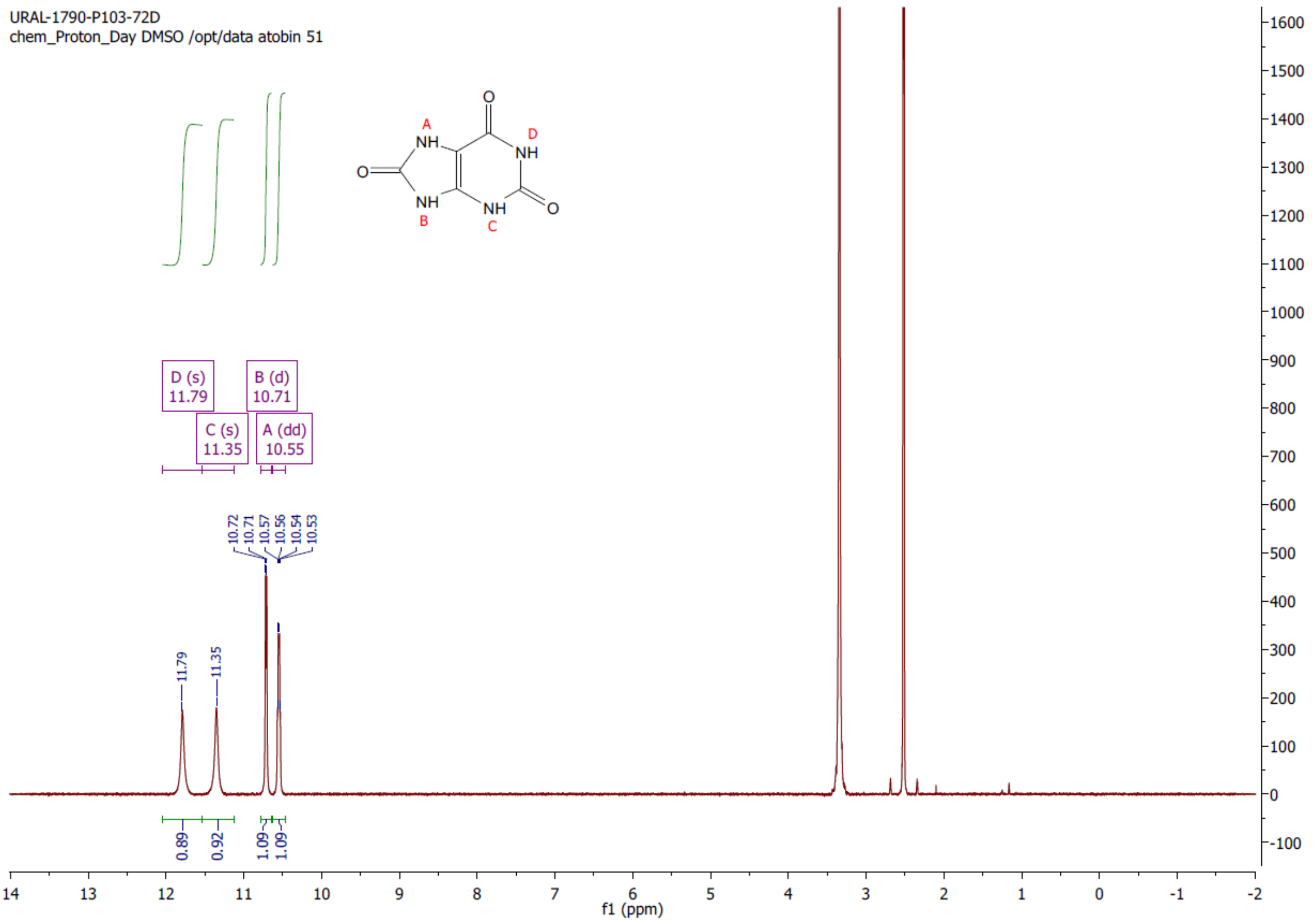

$^1$H-NMR Spectrum Conforms to Structure

Acanthus Research Inc.
2395 Speakman Drive, Suite 2111, Mississauga, Ontario, Canada L5K 1B3 Phone: (647) 478-1021
 www.acanthusresearch.com

### Analytical Summary

[13]C-NMR Spectrum

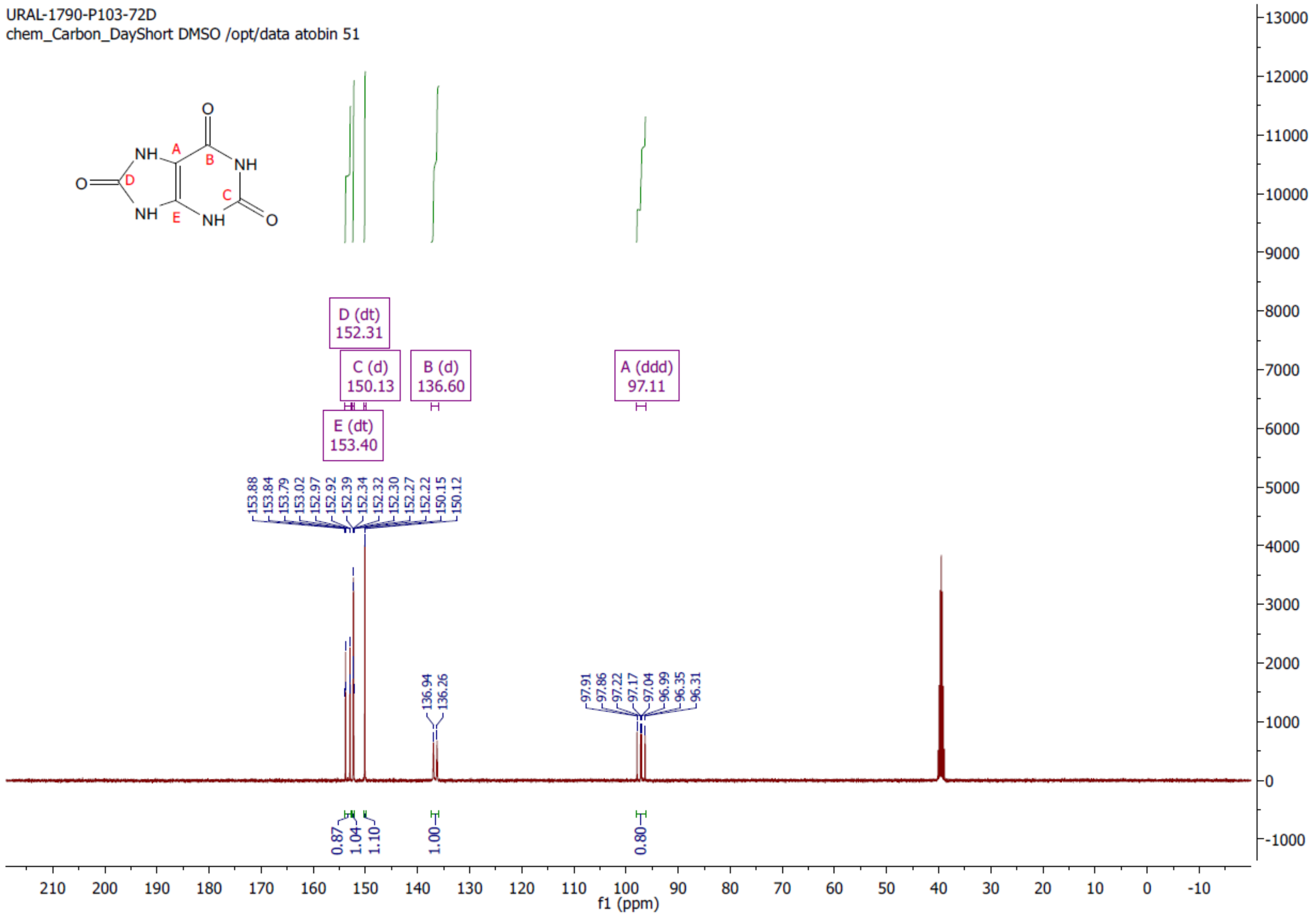

[13]C-NMR Spectrum Conforms to Structure

Acanthus Research Inc.
2395 Speakman Drive, Suite 2111, Mississauga, Ontario, Canada L5K 1B3 Phone: (647) 478-1021
 www.acanthusresearch.com

### Analytical Summary

| Mass Spectrum (ESI+) |
|---|

| | | | | | |
|---|---|---|---|---|---|
| **Sample Name** | URAL-1790-P103-72D | **Data File** | 211029_2094.d | **Acq Method** | HRMS.m |
| **DA Method** | AIMS_Accurate_Mass.m | **Instrument** | Agilent 6538 UHD | **Acq Date, Time** | 29-Oct-21 3:12:14 PM |
| **Comment** | ESI+ | | | | |

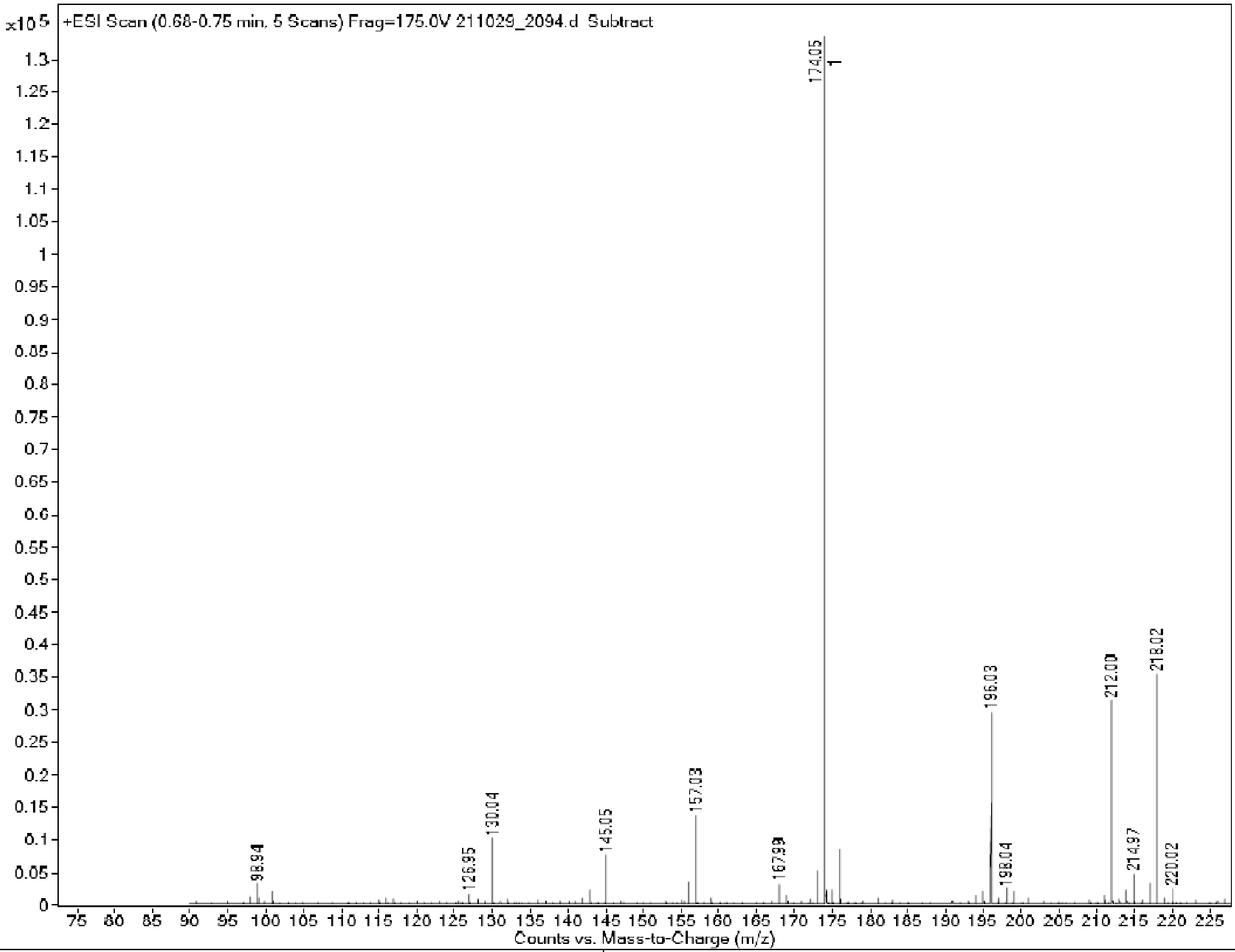

| | |
|---|---|
| Molecular Ion Peak $[M+H]^+$ | 174.05 |
| Mass Spectrum Conforms to Structure | |

Acanthus Research Inc.
2395 Speakman Drive, Suite 2111, Mississauga, Ontario, Canada L5K 1B3 Phone: (647) 478-1021
 www.acanthusresearch.com

### Analytical Summary

Mass Spectrum (ESI+) Expansion 1

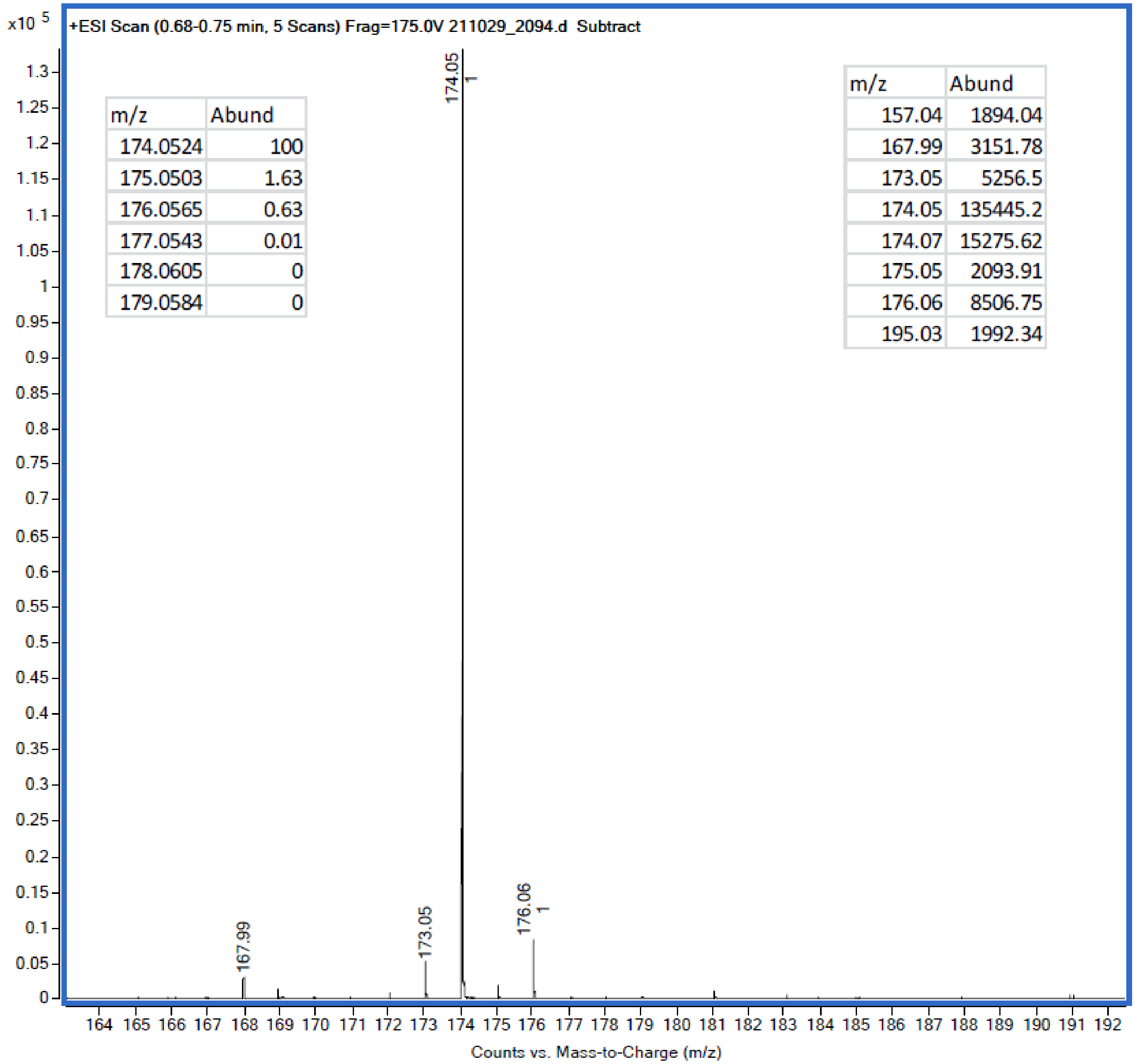

| m/z | Abund |
|---|---|
| 174.0524 | 100 |
| 175.0503 | 1.63 |
| 176.0565 | 0.63 |
| 177.0543 | 0.01 |
| 178.0605 | 0 |
| 179.0584 | 0 |

| m/z | Abund |
|---|---|
| 157.04 | 1894.04 |
| 167.99 | 3151.78 |
| 173.05 | 5256.5 |
| 174.05 | 135445.2 |
| 174.07 | 15275.62 |
| 175.05 | 2093.91 |
| 176.06 | 8506.75 |
| 195.03 | 1992.34 |

| | |
|---|---|
| Isotopic Purity | 99.3% |
| Unlabeled Content ($d_0$) | 0.0% |

Acanthus Research Inc.
2395 Speakman Drive, Suite 2111, Mississauga, Ontario, Canada L5K 1B3 Phone: (647) 478-1021
 www.acanthusresearch.com

### Analytical Summary

High Performance Liquid Chromatography

```
Acq. Operator   : Ryan Correa                    Seq. Line :   2
Sample Operator : Ryan Correa
Acq. Instrument : LC 6                            Location :   42
Injection Date  : 2021-10-29 9:54:33 AM                Inj :   1
                                                Inj Volume : 10.000 µl
```

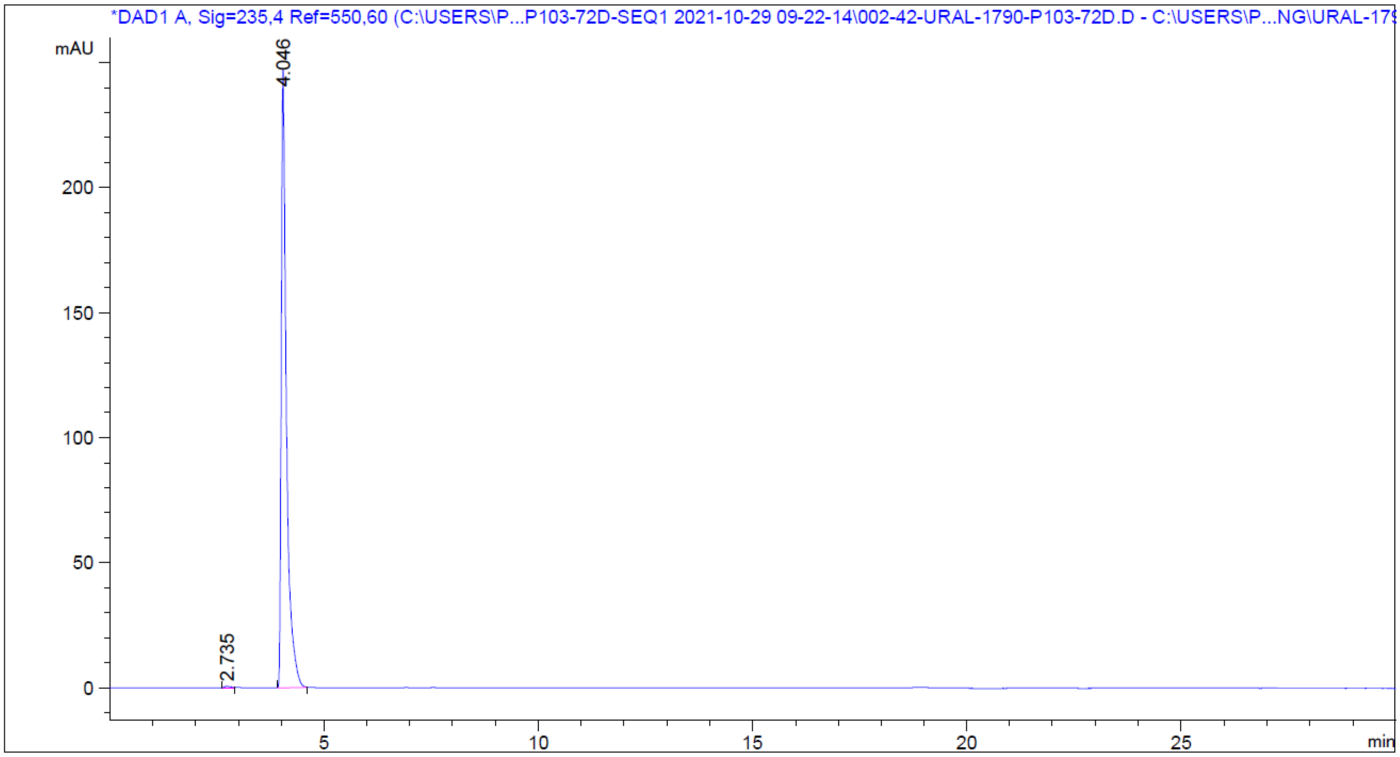

| Peak # | RetTime [min] | Type | Width [min] | Area [mAU*s] | Height [mAU] | Area % |
|---|---|---|---|---|---|---|
| 1 | 2.735 | BB | 0.1251 | 6.27090 | 7.38226e-1 | 0.2943 |
| 2 | 4.046 | BB | 0.1261 | 2124.51978 | 247.55750 | 99.7057 |
| Totals : | | | | 2130.79067 | 248.29572 | |

Chromatographic Purity 99.7%

Acanthus Research Inc.
2395 Speakman Drive, Suite 2111, Mississauga, Ontario, Canada L5K 1B3 Phone: (647) 478-1021
 www.acanthusresearch.com

### Certificate of Analysis

| | | |
|---|---|---|
| Product Number: | U-10826-01 | 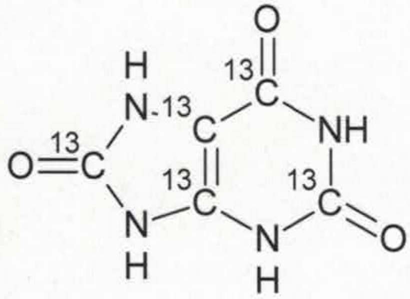 |
| Product Name: | Uric Acid-$^{13}C_5$ | |
| CAS No: | 69-93-2 (unlabeled) | |
| Molecular Formula: | $^{13}C_5H_4N_4O_3$ | |
| Molecular Weight: | 173.07 g $mol^{-1}$ | |
| Chemical Name: | 7,9-Dihydropurin-2,6,8-trione-$^{13}C_5$ | |

| | | |
|---|---|---|
| Lot Number: URAL-1790-P103-72D | Test Date: October 29, 2021 | Retest Date: October 2024 |
| Long Term Storage Conditions: Refrigerate | | |

| TEST | METHOD | RESULT |
|---|---|---|
| Appearance | Visual | Yellow Solid |
| Identification 1 | $^1H$ NMR | Conforms |
| Identification 2 | High Resolution Mass Spectrometry | m/z $[M+H]^+$<br>Theoretical: 174.0524<br>Measured: 174.0525 |
| Chromatographic Purity | HPLC; $\lambda$ = 235 nm | 99.7% |
| Isotopic Purity | High Resolution Mass Spectrometry | 99.3% (0.0% Unlabeled) |

| | |
|---|---|
| Use Statement: | For laboratory analytical use only; not for human consumption |

| | | | |
|---|---|---|---|
| Prepared by/Date:<br>(Analytical Services) | [signature] Nov 1 2021 | Approved by/Date:<br>(Quality Assurance) | Sw. Nov 1, 2021 |

Acanthus Research Inc.
2395 Speakman Drive, Suite 2111, Mississauga, Ontario, Canada L5K 1B3 Phone: (647) 478-1021
 www.acanthusresearch.com
